# Landscape structure and management alter the outcome of a pesticide ERA: evaluating impacts of endocrine disruption using the ALMaSS European Brown Hare model

**DOI:** 10.1101/025833

**Authors:** Chris J. Topping, Lars Dalby, Flemming Skov

**Affiliations:** Department of Bioscience, Aarhus University, Grenåvej 14, 8410 Rønde, Denmark

**Keywords:** Environmental Risk Assessment, Landscape Generation, Agricultural System Simulation, Pesticide Regulation, Abundance Occupancy Ratio

## Abstract

There is a gradual change towards explicitly considering landscapes in regulatory risk assessment. To realise the objective of developing representative scenarios for risk assessment it is necessary to know how detailed a landscape representation is needed to generate a realistic risk assessment, and indeed how to generate such landscapes. This paper evaluates the contribution of landscape and farming components to a model based risk assessment of a fictitious endocrine disruptor on hares. In addition, we present methods and code examples for generation of landscape structures and farming simulation from data collected primarily for EU agricultural subsidy support and GIS map data.

Ten different Danish landscapes were generated and the ERA carried out for each landscape using two different assumed toxicities. The results showed negative impacts in all cases, but the extent and form in terms of impacts on abundance or occupancy differed greatly between landscapes. A meta-model was created, predicting impact from landscape and farming characteristics. Scenarios based on all combinations of farming and landscape for five landscapes representing extreme and middle impacts were created. The meta-models developed from the 10 real landscapes failed to predict impacts for these 25 scenarios. Landscape, farming, and the emergent density of hares all influenced the results of the risk assessment considerably.

The study indicates that prediction of a reasonable worst case scenario is difficult from structural, farming or population metrics; rather the emergent properties generated from interactions between landscape, management and ecology are needed. Meta-modelling may also fail to predict impacts, even when restricting inputs to combinations of those used to create the model. Future ERA may therefore need to make use of multiple scenarios representing a wide range of conditions to avoid locally unacceptable risks. This approach could now be feasible Europe wide given the landscape generation methods presented.

## 1. INTRODUCTION

In Europe pesticides are regulated under Regulation (EC) 1107/2009, a replacement for Directive 91/414/EEC. This change in regulation has sparked a gradual change the regulatory focus from individual toxicity and single compound regulation towards a population approach, and consideration of ecosystem services as a key component of regulatory environmental risk assesssments (Nienstedt, Brock et al. 2012). In the USA, similar considerations have been applied to environmental risk assessment (ERA), and landscape scale ERA has already been undertaken (Landis 2003). In the EU this step has been proposed recently by EFSA (EFSA Panel on Plant Protection Products and their Residues (PPR) 2015, Topping, Craig et al. 2015). This change towards landscape scale requires consideration of many new facets of ecology in the ERA, very different from the traditional hazard quotient and toxicity-exposure ratio approach. Although it seems there is a general concensus that population models have the potential for adding value to ERA by incorporating better understanding of the links between individual responses and population size and structure and by incorporating greater levels of ecological realism, there are still many issues that require further study (Forbes, Calow et al. 2008).

One of the new issues that need to be dealt with is the fact that the precise effect of pesticide application in a particular landscape configuration relies on complex spatial and temporal dynamics involved in animal behaviour and ecology. For example, in ERA focus is often placed on recovery of local populations, utilizing the spatial dynamics of mobile agricultural land species. However, this recovery is normally based on small plot experiments which do not take into account the landscape scale impacts of source-sink dynamics (Topping and Lagisz 2012, Topping, Kjær et al. 2013, Focks, ter Horst et al. 2014). In addition, the way in which the pesticide is used, and the context into which it is placed is also likely to alter population outcomes (e.g. 20% by area treated in a predominatly arable area may have a different effect to the same area in a pastoral context).

In conservation and general ecology it is well known that landscape structure will influence population dynamics, and population persistence (e.g. Silver, Wooster et al. 2004, Aviron, Kindlmann et al. 2007, Pavlacky, Possingham et al. 2012), and as a consequence that it can affect biological control in agro ecosystems (With, Pavuk et al. 2002). However, until recently there was little focus on the role of landscape structure in determining impacts in ERA. In some cases landscape features have been used to more precisely determine predicted exposure (Gaines, Porter et al. 2005), or as an explanatory variable in estimating population impacts, but rarely taken into account as a determining factor for the state of the population before exposure to pesticides. However, previous simulations using voles and carabid beetles have shown that precise structure is important to predict pesticide impacts, i.e. both composition and configuration of landscape elements must be considered (Dalkvist, Sibly et al. 2013, Topping, Craig et al. 2015). Similarly it can be expected that the context of the pesticide application will also be important if it changes the properties of the population in focus. This will include both landscape structure and management contexts altering exposure as well as potentially the state of the organisms pre-exposure.

Recent EFSA scientific opinions (EFSA Panel on Plant Protection Products and their Residues (PPR) 2015, EFSA Panel on Plant Protection Products and their Residues (PPR) 2015) highlight the need for landscape approaches, and a number of studies include spatial issues in pesticide ERA (e.g. Topping and Lagisz 2012, Liu, Sibly et al. 2013, Meli, Auclerc et al. 2013). However, many approaches utilize very simplified representations of the spatial structure of the landscape.

Landscapes represented by one or a few regularly shaped fields are common, despite the fact that when linked to population models it is known that these regular representations result in bias (Holland, Aegerter et al. 2007). There must therefore also be an open question regarding neutral landscapes (With 1997) generated by more complex artificial procedures. In these landscapes, statistical structure may be comparable, but functional structure may not be maintained (e.g. roadside verges by roads, field geometry and field margin placement).

There may be four reasons for not using realistic landscapes. The first is an active choice of a simple representation for simplicities sake. The second may be that computational limits resulting from hardware or language limitations restrict the complexity of the system modelled. The third may be that creation of sufficiently accurate and detailed landscape representations is difficult in itself. Fourth, if it was demonstrated that landscape realism has no impact on the ERA. In this study, we address the question of the extent of the impact precise landscape and farming configurations have on an ERA. This is done using a range of Danish landscapes and their associated management created from EU farming subsidy claim data together with readily available GIS mapping data.

To be able to make a realistic assessment of landscape and management impacts it was necessary to be able to create realistic maps and farm management. The method for creating landscapes which can be used as a basis for landscape-scale population-level risk assessment is presented, as well as details of the ALMaSS farm management module. The landscape generation method involves the linking of a number of data sets not previously possible due to data restrictions. It is therefore a new a powerful procedure that may be widely applicable.

This paper considers how landscape structure and management affects ERA by carrying out what might be considered a higher tier ERA of a fictitious endocrine disruptor by modelling hare populations in various landscape scenarios in ALMaSS (Topping, Hansen et al. 2003). It is important to note that the focus of this paper is the interaction between landscape features and ecology and behaviour of the hare on an ERA. It is not the intention to evaluate a specific mode of action or pesticide, but to provide proof of concept for the need to consider these interactions in ERA. Hence, we have chosen a particular scenario, but do not suggest it to be the only useful toxicological situation to consider, or that this case is representative of all others. As a consequence we do not go into all possible toxicological details.

## 2. METHODS

This paper present a simulation of pesticide application scenarios in ten real Danish landscapes. The simulation models used are part of ALMaSS (Topping, Hansen et al. 2003), a large simulation system comprised of many interacting agent-based models. ALMaSS has two main components: the dynamic environment or landscape that change over time as a result of farm management and season and the animal agents that are affected by the landscape and interact with other agents. This section is divided in three: The making of the landscape is described in *“Landscape generation”* below and the agent-based modelling is treated under *“ALMaSS simulation system”.* Finally, the choice of pesticide application scenarios and analyses is described under *“Scenarios”.*

### 2.1 LANDSCAPE GENERATION

All landscapes used in the current study are surface covering raster maps with a resolution of 1 m * 1 m. They were all generated using the same mapping algorithm and input data sets. The overall process consists of the following six steps:

1. Classifying and defining farm types;
2. Stratified selection of model landscapes;
3. Converting input vector data to raster layers;
4. Combining individual raster layers into thematic maps (e.g., all road types, paths and railway tracks in a transportation theme);
5. Stacking thematic maps in a reasonable order (i.e. roads on top of fields etc.), and, finally;
6. Reclassifying and regionalising the land cover map to an ALMaSS landscape.

Below we first describe the input data, then the steps involved in generating the final map. For further details see Appendix 1 and the program code (in Python) available at: https://github.com/au-bios/python-landscapegen.

#### 2.1.1 INPUT DATA

We used publicly available vector map layers from the Danish common public geographical administration data (GeoDanmark data, downloaded 2012, http://download.kortforsyningen.dk). The vector layers were used to map 41 different landscape features (see Table S1).

The Danish AgriFish Agency (under the Ministry of Food, Agriculture and Fisheries) maintains an annually updated map of all fields and a database of crops grown in Denmark (“Det Generelle Landbrugsregister” - GLR) (Danish Ministry of Food Agriculture and Fisheries 1999). Farmers are obliged to report for each individual field the crop they intend to grow the following year. The data set used in this study is from 2013 where more than 45,000 farmers contributed to the database. The data set makes it possible to identify the owner (or manager) of each field and the actual crop grown on it.

Lastly, where a pixel in the final land-cover map was not covered by either a field or any of the GeoDanmark layers (approximately 3-5% of the area), we used the Area Information System (AIS) data, which is a surface covering map with 45 different land cover types for Denmark (Stjernholm, Olsen et al. 2000). The AIS map is based on data and satellite imagery from the late nineties, but is, nevertheless, the best surface covering map available.

Soil types, used for farming and vegetation growth purposes in ALMaSS, are also mapped. To map soil type to each field in the landscapes we used a soil classification map (resolution 1:200.000) from the Danish centre for food and agriculture (DCA) at Aarhus University (downloaded from http://dca.au.dk/forskning/den-danske-jordklassificering/). The soil classification map was rasterized and field polygons were overlaid to determine the dominant soil type for each field.

During an ALMaSS simulation, crop type on any field at any time is a function of management (crop rotation based on the GLR data), weather and the soil type of the field. The crop rotation also depends on the farm type, which is created by classifying GLR data together with data on the numbers and type of stock a farm has. This data is available from the Danish Livestock Register (CHR) (Danish Ministry of Food Agriculture and Fisheries 1999) a data set used primarily for disease control in Denmark.

#### 2.1.2 STEP 1: FARM CLASSIFICATION AND ROTATION

A small program was used to classify all farms in Denmark into general farm types (see Appendix 1 for documentation and links to program code and its documentation). The program classifies all farms based on a combination of the crops they are growing using data obtained from the GLR, and on the animals they have from the CHR.

By combining crop and animal information, it was possible to identify major farm types such as pig or arable, or dairy farms. Some less common types are also identifiable, e.g. farmers that grow sugar beet on contract. In addition to this information the GLR also indicates whether a farm is organic or not and the overall farm size. This extra information provides the basis for the classification. Rules used to classify the farms needed to be very general because real farms tend not to fit neatly into pure farm type rules (e.g., many arable farms have grazing because they have some animals e.g. horses or a few animals for their own consumption). The rules used were:

1. Farms with large proportion of vegetables (minimum 0.5) and larger than 2-ha were organic or conventional **Vegetable** farms, otherwise if small were classified as ‘other’.
2. Farms with a proportion of potatoes or sugar beet not less than 20% were **Potato** or **Beet** farms respectively.
3. Farms with animal (cows, sheep and pigs) transformed to standard animal units that have fewer than 20 animal units and an area less than 20 ha were designated as **Hobby** farms (<25 ha is typically part-time or hobby (Levin 2006), so 20 ha will reduce the chance of misclassifying commercial farms).
4. Farms with animal units above 20 and cattle + sheep above 75% they were designated as **Cattle** farms.
5. Farms with animal units above 20 and pigs above 75%, or crop area of grazing pigs above 15% were designated as **Pig** farms.
6. Farms with animal units above 20, but not pig or cattle farms, were designated as **Mixed Stock**.
7. Farms with no animals registered but with large areas of grazing were assumed to be either **Cattle** farms or **Mixed Stock** farms depending on whether grazing area was above 40% or 20-40% respectively.
8. All remaining farms must have been **Arable** farms (i.e. large area with few or no animals and little or no grazing).

All farms except ‘other’ could be designated as organic or not dependent upon the information on that farm in the GLR giving a total of 17 farm types possible.

For each farm type, the mean proportion of the farm crop area was calculated for each crop. Crops with less than 1% share of the area of a farm type were ignored and the rest used to create a farm rotation for that classification. It was assumed that the rotation could be represented by 100 crops (1 crop for each 1%). The order of crops followed typical agronomic practices and issues such as late harvest leading to impossible sowing conditions were controlled by the built in ALMaSS farm code. The result is a pattern of changing crops on a field that matches the overall crop distribution pattern for that farm type precisely over 100 seasons. Viewed on a larger scale crop distributions will therefore be overall correct at any point in time, although the actual crop grown on a single field will not replicate reality. This method does not, however, take into account differing soil types between fields, which in reality would restrict some crop distributions.

#### 2.1.3 STEP 2: LANDSCAPE SELECTION

We created ten landscapes representing different predominant farming/landscape structure combinations for use in the pesticide scenarios. Initially all farms were classified and mapped at national scale. ‘Hot-spots’ where particular farm types dominated were identified and the landscapes selected to capture as much variation as possible whilst having all main types represented. The ten 10 x 10 km landscapes cover predominantly agricultural areas without large woodlands or towns. Predominant farm types were conventional pig (2 landscapes), conventional cattle (3), conventional arable (2), conventional potato (1) conventional sugar beet (1) and organic cattle (1) (see Appendix 3).

#### 2.1.4 STEP 3: CONVERSION OF VECTOR- TO RASTER-LAYERS

Each of the vector layers were initially rasterized with a resolution of 1 m * 1 m. Linear features were described as their centre line. To get a meaningful raster representation of these features, a buffer was added when converting to raster. For example for large roads (width 10m) we added a 5m buffer around the centre line. Same procedure was followed for other linear features (e.g. streams and hedgerows). Point features such as wind turbines, power pylons and individual trees which are represented as their centre points were also buffered (see the online code for details). Fields are a special case since identification of individual fields is necessary for the farm management in ALMaSS.

When converting the vector layer of all fields, each field was assigned a unique id. All input data initially used the UTM zone 32/ETRS89 coordinate system.

#### 2.1.5 STEP 4: COMBINING INDIVIDUAL RASTER LAYERS INTO THEMATIC MAPS

Individual layers were organized into thematic maps before combining to the final map. E.g. all layers with roads, road verges, road side slopes, tracks, railroads etc. were combined in to a road theme. In total the 41 layers were combined into six different themes (built up areas, nature, wet nature, water and cultural areas). At this point, each theme is a raster map with the value 1 where features in the theme are absent or another numeric value unique to the theme where the features are present.

#### 2.1.6: STEP 5: STACKING OF THEMATIC MAPS

Finally the themes were stacked step by step onto each other. Since each data source has variable accuracy, and some features are overlaid, a procedure was needed to ensure complete coverage, this was based on 9 layers and maps:

- The field layer was used as the bottom layer (1)
- The lake and streams were put on top of the field layer (2)
- Dry natural areas were then added, but only to cells not already occupied by 1 or 2 (3)
- Built-up areas were added to cells not already occupied by 1, 2 or 3 (4)
- Wet natural areas (5)
- Cultural features (6)
- Roads (7)
- Sea (8)
- Buildings (9)

Layers 5-9 were added sequentially on top of everything else.

After this process there may still be a number of cells without land cover type, depending of the quality of the input data. For the present study we used an existing, older land cover map of Denmark (AIS data) to fill in these gaps.

All handling and analysis of spatial data was done using Python and the python library *arcpy* to access ArcGIS features. The python script used to generate the maps used here is freely available on Github (https://github.com/au-bios/python-landscapegen).

#### 2.1.7 STEP 6: RECLASSIFY AND REGIONALIZE LAND COVER MAPS

The land cover map contains more detail than are used in ALMaSS for the purpose of this study and need to be condensed into the landscape element types to be used in the simulation system. This was done using a simple reclassification of sub-divided types to ALMaSS landscape element types. Finally, as the landscape simulation engine in ALMaSS works on polygons rather than individual cells, clusters of cells with similar land cover class were converted into polygons using regionalizing and exported in ASCII raster format with each cell of the raster containing the polygon reference number for the polygon occupying the majority of that 1m^2^. ALMaSS therefore uses the flyweight pattern (Gamma, Helm et al. 1994) with each cell in the raster containing a reference number of a polygon where all data concerning that polygon is stored. Polygons are considered to be homogenous.

### 2.2 ALMaSS SIMULATION SYSTEM

For a comprehensive description of the ALMaSS, the reader is directed to the online documentation found at (Anon. 2014). This documentation follows ODdox format (Topping, Hoye et al. 2010), combining model description with doxygen (van Heesch 1997) code documentation.

ALMaSS is comprised of two main components, the environment and its associated classes and the animal representations (classes). The environment interface is provided by the ‘Landscape’ class. This class contains a map of the landscape to be simulated together with individual landscape elements such as fields, hedges, roads and woodlands. Fields are a special case. Fields are linked in groups to form farms. These groups were based on ownership or management information from municipal or EU-farming subsidy sources (see Landscape Generation). Each farm is an instance of the Farm class which simulates the detailed management of its fields, dependent upon its farm type, the weather, soil type, and past history of management.

Crop management is an important feature of ALMaSS. Although originally implemented in 2003 (Topping, Hansen et al. 2003), ALMaSS still appears to be unique in being able to generate dynamic patterns of crop usage at landscape scales together with detailed crop husbandry. This produces emergent properties of crop distribution between fields and proportions of crops at the landscape scale, but also of patterns of crop management activities in space and time (e.g. see Topping and Odderskær 2004, Gevers, Hoye et al. 2011, Parry, Topping et al. 2013). The farm management and crop rotation methods in ALMaSS are documented previously (Topping 2009) but a written description has also been provided in Appendix 2.

All vegetated landscape elements (crops and non-crops) undergo type-specific daily vegetation development based on weather and fertilizer inputs as drivers. Farm management events (e.g. spraying, harvest, ploughing) directly interact with vegetation height and biomass, providing a dynamic picture of changing landscape conditions as a result of both environmental and anthropogenic processes and factors. These events may impact animals directly (e.g. ploughing related mortality) or indirectly e.g. pesticide application resulting in residues and subsequent exposure.

The second main ALMaSS component is the simulation of animals, represented by specific species classes all derived from a common base class. All animals are agents and are affected by environmental variables, vegetation structure, and by direct interaction with other agents or farm management. Each animal represents an individual of a particular species, with its own behavioural rules and interactions with its environment. Animals can sense the characteristics of their environment (habitat type, vegetation structure, temperature etc.), management events, and their own physiological condition. Hence, animals exposed to management will choose behaviour suitable for that management, their current location, and physiological state. Animals can interact with each other in a variety of ways ranging from simple local-density dependent interactions to complex behavioural messaging, depending upon animal type and current activity. All animals share a common basic form of control simulated as a state machine. This means that they exhibit behaviour associated with a specific state, and make transitions to other behavioural states as a result of internal or external cues.

#### 2.2.1 HARE MODELLING

The animal model used for all simulations here was the European Brown Hare model (Topping, Hoye et al. 2010). The model simulates the growth, movement, reproduction, and mortality of individual hares using a daily time-step for most activities, but a 1 minute time-step for foraging. Full details of the hare model are described by (Topping 2009), but a short description is presented here to aid readability.

The hare model simulates five life-stages: *infants* up to 11 days during which they are totally dependent on the lactating doe; *young* 12-35 days old after which they are fully weaned, *juveniles* 35-365 days old, adult *males* and *females.* In the model hares are quite mobile and able to find suitable forage over a wide area when not encumbered with young. If feeding conditions are good hares will generally drift over large areas, in poor feeding conditions the hares will optimally forage, and thus may become restricted to localised ranges for periods within a season. Breeding starts in spring if body condition allows for the production of foetal mass. After birth the female must increase her energy intake in order to provide enough energy for lactation. Energy comes from foraging from green shoot material and the amount of energy obtained depends on the age of the shoot and the overall structure of the vegetation. Dense vegetation may therefore have a high food value in terms of biomass but a poor digestibility and high impedance. A female that cannot support lactation because her combined energy intake and reserves fall too low will abandon her young. Reproduction will not be attempted again until energy reserves are replenished. Growth of model hares is also dependent on energy balance and hares which do not achieve 45% of their potential weight at any age will die. Adults rarely die of energy shortage and are assumed to be able to “carry” a negative energy balance. They thus will remain in the population contributing to social stress, which is ultimately the primary density-dependent regulation factor in the hare model and reducing population growth. Hunting occurs in autumn but other non-energetic related losses are based on life-stage specific constant daily probabilities (e.g. predation of young), or on events driven by human management activities (e.g. harvest mortality).

#### 2.2.2 PESTICIDE MODELLING AND TOXICOLOGY

The model hares must forage realistically from the landscape and are exposed to pesticide residues in the process. Foraging is done by selecting a 10 × 10 m area at the current hare location and foraging from this; this is assumed to take 100 minutes. The hare then selects the 10 × 10 m area adjacent to this first forage area that provides the best energetic return based on forage quality and impedance. Time to walk between squares and sample them is also included. This process continues until either the period of time allocated for foraging is used up, or the hares cannot eat any more.

Hares have an energetic maximum daily intake limit of 5500 kJ day^-1^ (Valencak, Tataruch et al. 2009), but stomach contents of wild hares contain 11 kJ g^-1^ (Hacklander, Tataruch et al. 2002), which suggests a daily throughput maximum of 500 g. The maximum rate observed for hares by Andersen (1947) was 1.7g per minute would result in only 294 minutes foraging. This is much less than assumed in ALMaSS where 67% of the daily activity will be foraging or movement associated with foraging. To compensate we assume that intake rate in grams per minute is 500/(1440 min × 0.67) = 0.518 g per foraging minute. If the hare uses less time than this due to other activities, then the pesticide intake rate will decrease proportionally with the forage intake. The ingestion rate of pesticide (mg minute^-1^) is therefore the environmental concentration (mg/g) multiplied by 0.518 for each minute spent foraging in each location.

The model includes internal and external toxicokinetics (TK) in terms of the varying rates of ingestion of the pesticide, and the process of elimination within the hare. The internal TK are represented by a single compartment model assuming a percentage elimination rate per day. External TK is determined by the feeding behaviour of the hare and ultimately by the time spent feeding from contaminated areas, and the concentration of pesticide on vegetation.

We based the scenarios on a generalised pesticide. Application rates are assumed to be 10 g a.i. per hectare. This gives a residue of 0.4 mg/kg vegetation immediately after spraying based on a mean residual unit dose (RUD) for cereals and leafy forage crops (Fletcher, Nellessen et al. 1994). This will result in a daily dose of 0.2 mg a.i. per day (0.5 kg/d × 0.4 mg/kg = 0.2 mg/d), if an adult hare eats its full 500 g from a contaminated area immediately after spraying. Per foraging minute we can calculate that the initial rate after spraying would be 0.000518kg * 0.4 mg/kg = 0.000207 mg/min. We assume an environmental half-life of 7 days.

Drift was included, assuming that drift occurs up to 12 m from the edge of any sprayed field, following the equation *p* = *e*^−0.6122^*^x^*, where *p* is the proportion of application rate falling from x to 1m, and x is distance in m from the point of spray. This gives c.a. 24% drift at 1 m, and 2.1% at 5 m. The direction of drift varies randomly depending on the day of spraying and was assumed to be due north, south, east or west.

In all cases it was assumed that the pesticide was an insecticide. Spraying on spring barley harvested at maturity (i.e. not spring barley for silage), was with a 35% chance of one application each season. For winter wheat grown to maturity there were up to three applications per season each with a 50% chance dependent on carrying out the previous spray (3 applications probability is therefore 12.5%). Spring barley application was on April 30^th^, winter wheat applications on 15^th^ May, 1^st^ June and 15^th^ June. No other crops were assumed to be treated with toxic substances, although otherwise followed normal agricultural management. These treatment probabilities allowed calculation of mean frequency of application per landscape by calculating the area of barley and wheat and multiplying by their respective treatment frequencies.

The toxicological effects assumed were entirely constructed and are not intended to represent any real pesticide, but were designed to demonstrate noticeable impacts. The effect used was intended to represent a chronic reproductive effect. Change in litter size is typical of such an impact for pesticides on mammals (Bishop, Morris et al. 1997, Sharara, Seifer et al. 1998), and we assumed that this was the only impact (typically litter number and offspring size can also be affected). For the chronic effect, the impact of exposure above a threshold body-burden is modelled as a uniformly distributed chance of litter size reduction of 0-100% for female hares exposed during gestation of that litter. Initial scoping runs indicated that a trigger threshold of 0.0001 mg a.i./kg bw with an internal degradation rate of 5% per day gave noticeable population level impacts. This was designated as the 1X toxicity scenario. Increasing sensitivity by factor 10 to 0.00001 mg a.i./kg bw was designated the 10X toxicity scenario.

### 2.3 SCENARIOS

Ten landscapes of 10 × 10km area were selected using the landscape generation methods described above. These landscapes varied in structure and had differing farm management based on differing predominant farm types (Pig (2 landscapes), Dairy (3), Arable (2), Organic Dairy (1), Potato (1) and Sugar beet (1)). They were also chosen to span the range of hare densities encountered in Denmark, from high (Lolland, sugar beet) to low (Mors, pig farms) (Table 1). See Appendix 3 for a figure of each landscape and Appendix 4 for the proportion of area assumed occupied for each crop for each farm type. The proportion of the area of landscape occupied by each main crop and the total area treated with pesticide varied between landscapes. Treated area varied from 35% to 60% of the farmed landscape (Table 2).

**Table 1.** Ten landscapes (Esbjerg – Tønder) with the for most common farm types by area listed. Conv. = conventional. Org. = organic.

| Esbjerg |  | Himmerland |  | Karup |  | Kolding |  | Lolland |  |
| --- | --- | --- | --- | --- | --- | --- | --- | --- | --- |
| Type | %Area | Type | %Area | Type | %Area | Type | %Area | Type | %Area |
| Conv. Cattle | 75% | Conv. Cattle | 65% | Conv. Potato | 84% | Conv. Arable | 59% | Conv. Beet | 86% |
| Conv. Arable | 10% | Conv. Pig | 11% | Conv. Cattle | 6% | Conv. Pig | 20% | Conv. Arable | 11% |
| Conv. Mixed Stock | 4% | Conv. Mixed Stock | 9% | Org. Cattle | 3% | Conv. Cattle | 12% | Conv. Mixed Stock | 1% |
| Conv. Pig | 4% | Conv. Arable | 8% | Conv. Hobby | 2% | Conv. Hobby | 4% | Conv. Cattle | 1% |

| Mors |  | Næstved |  | Odder |  | Toftlund |  | Tønder |  |
| --- | --- | --- | --- | --- | --- | --- | --- | --- | --- |
| Type | %Area | Type | %Area | Type | %Area | Type | %Area | Type | %Area |
| Conv. Pig | 56% | Conv. Arable | 61% | Conv. Pig | 42% | Conv. Cattle | 39% | Conv. Cattle | 60% |
| Conv. Cattle | 18% | Conv. Mixed Stock | 16% | Conv. Cattle | 14% | Org. Cattle | 26% | Org. Cattle | 17% |
| Conv. Arable | 18% | Conv. Pig | 8% | Conv. Arable | 36% | Conv. Arable | 14% | Conv. Arable | 13% |
| Conv. Hobby | 5% | Conv. Cattle | 7% | Con. Hobby | 4% | Conv. Pig | 10% | Conv. Pig | 5% |

**Table 2.**
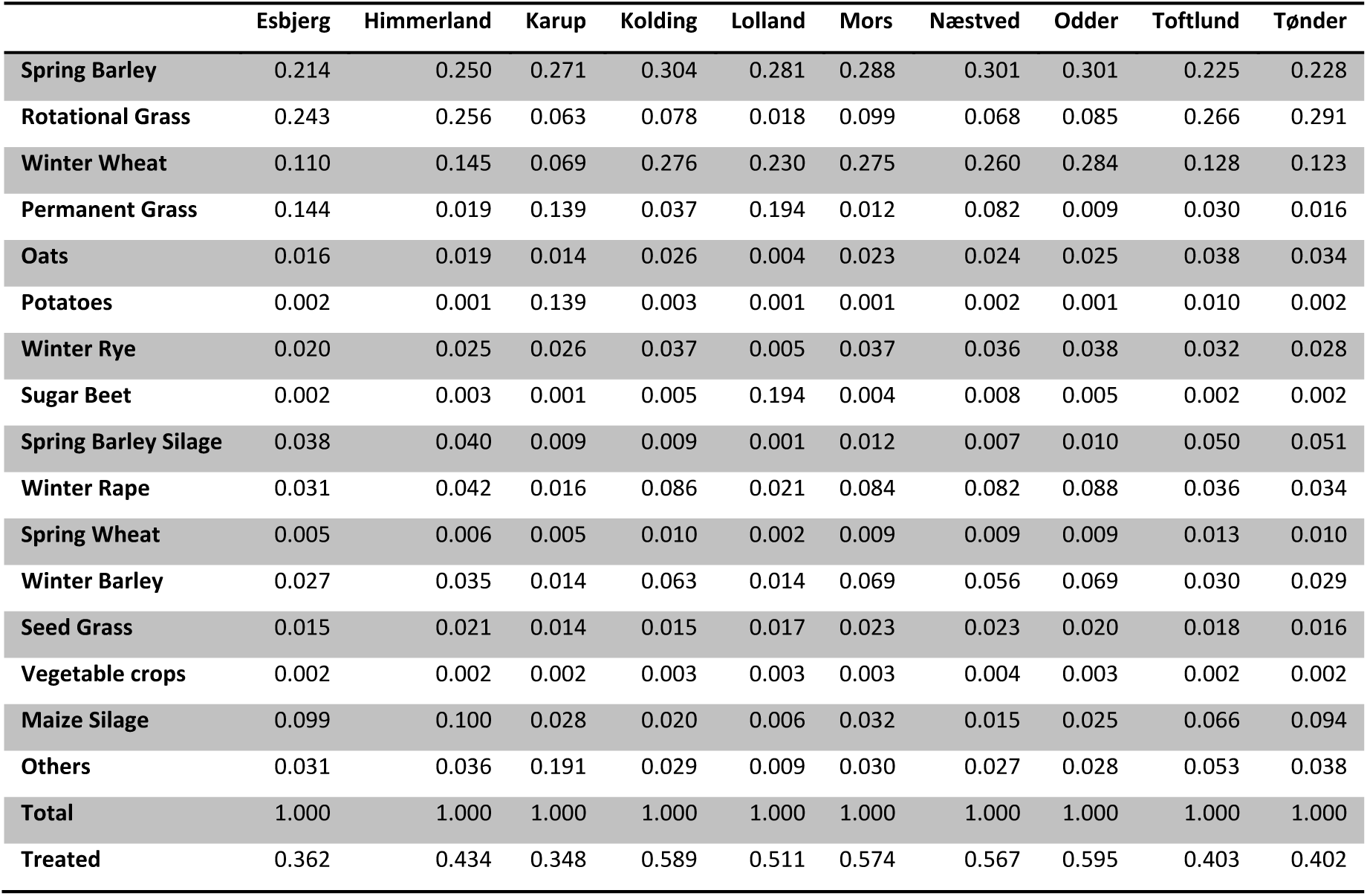
The proportion of each ALMaSS crop, ordered by overall area, by area in each of the ten landscapes used for the study. ‘Treated’ is the sum of the area treated with the pesticide for which the ERA was carried out (winter wheat and spring barley).

We considered four types of scenario for each:

1. Baseline scenarios using the correct farm classification and rotation for each landscape;
2. A treatment scenario using an application of our test pesticide using both acute and chronic toxicological impacts separately for each landscape using both 1X and 10X toxicity;
3. Baseline scenarios using farming information from another landscape;
4. Treatment scenarios using baseline conditions from ‘3’ and an application of our test pesticide using both acute and chronic toxicological impacts separately for each landscape using 1X toxicity.

Scenarios 3 & 4 were used to evaluate landscape and farming independently. These scenarios used farm management combinations from Esbjerg, Karup, Lolland, Odder and Tønder, as representing the range of density options from the baselines. In all cases all 20 replicates were run with all five farm classifications (an additional 20 combinations in addition to the five baselines).

For all scenarios simulations were run for 30 years and the pesticide was applied in the last 10 years (21-30).

### 2.4 ANALYSIS

Each scenario was run by running 20 replicates and analysed separately before creating an average response statistic of the 20 replicates. These response statistics were used to compare impacts. We used the Abundance Occupancy Ratio index (AOR-index (Hoye, Skov et al. 2012)) to compare responses between scenarios using a fixed grid size of 400m. The AOR-index provides a measure of changes in occupancy and abundance of hares relative to a baseline scenario. If we assume equal weighting to both changes in occupancy and abundance, then a summed AOR score can be made by summing the changes in both dimensions. This provides a reasonable guide to overall impact and is more sensitive than using the more typically employed measurement endpoint of population size (e.g. Dalkvist, Sibly et al. 2013).

If using AOR scores for ERA then typically we are concerned with a specific protection goal (SPG), which if described at the population level might be ‘negligible effects’ (see e.g. EFSA Panel on Plant Protection Products and their Residues (PPR) 2015). This could be done using the AOR approach to integrate both spatial and temporal impacts. In this case a SPG would need to be specified in terms of population impacts (e.g. a trigger value of 10%); then any scenario resulting in a summed AOR score of < -0.1 would trigger concern (see Fig. 1 for a graphical representation).

**Figure 1.**
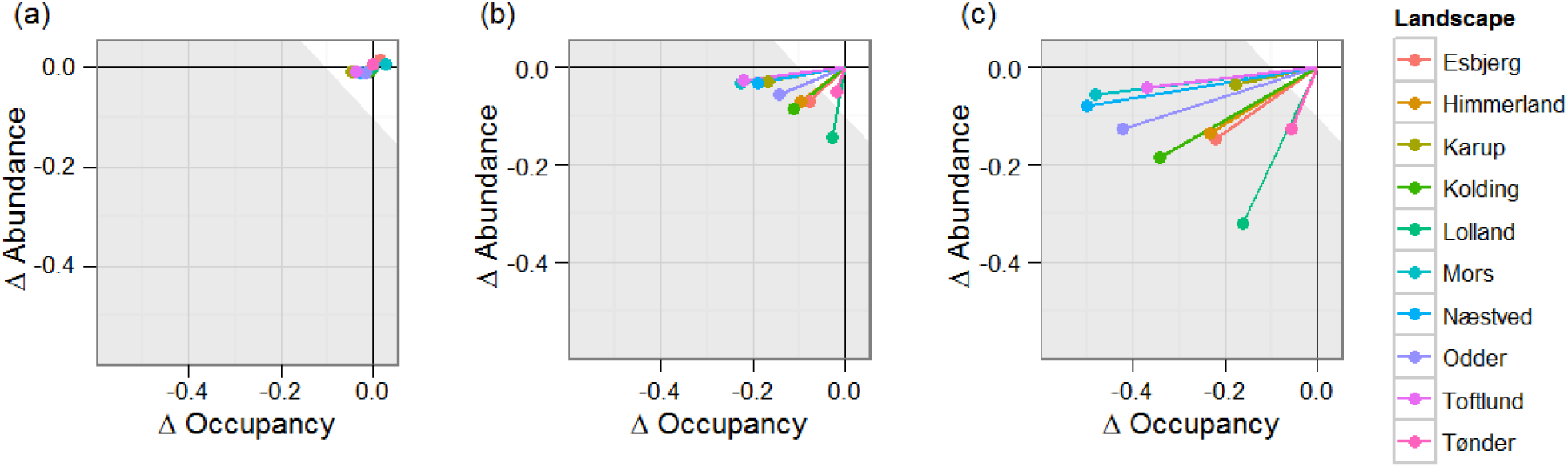
AOR scores from endocrine disruptor scenarios. a) AOR scores for the 10 different simulation landscapes relative to baselines in the 10 years prior to pesticide treatment for 10X toxicity. These scores can be viewed as the residual variation that cannot be attributed to the treatment in the following 10 years; b) AOR scores for the same landscapes, as a mean of the 10 years of pesticide treatment for 1X toxicity. c) The AOR scores for the 10X toxicity scenario. a-c) The white area exemplifies a SPG for a threshold of effect, assuming a SPG trigger value of -0.1 summed AOR score.

## 3. RESULTS

Baseline hare densities were predicted to vary between the landscapes (Table 3). Variations in density match the estimates for differences between landscapes reported for Denmark to date (ref Hare Management Plan). Densities on two places from the Lolland region were 65 and 111 hares km^−2^. Other regions varied between 3.1 and 13.0 hares km^−2^, 13 being a count from the Tønder area. Thus, although not possible to directly test the model against data from precisely replicated conditions, the general trends and range of densities seems to match those recorded.

**Table 3.** Baseline hare densities predicted for each landscape. Es = Esbjerg, Hi = Himmerland, Ka = Karup, Ko = Kolding, Lo = Lolland, Mo = Mors, Na = Næstved, Od = Odder, To = Toftlund, Tø = Tønder

| Landscape | Es | Hi | Ka | Ko | Lo | Mo | Na | Od | To | Tø |
| --- | --- | --- | --- | --- | --- | --- | --- | --- | --- | --- |
| Total Hares/km<sup>2</sup> | 11.31 | 11.97 | 2.94 | 21.15 | 65.24 | 3.72 | 7.94 | 2.17 | 1.25 | 28.50 |

Evaluating the impact of the endocrine disruptor at 1X and 10X toxicity using AOR scores was performed by comparison of mean AOR score over the last 10-years of simulation. In order to be sure that the deviations from the baseline were significant, and not due to between run variation, the analysis was performed for the 10 simulation years before pesticide application. The variation here can be considered to be the range of variation which cannot be attributed to the treatment differences. This variation was largest in the 10x toxicity replicates (Fig. 1a). This background variation is clearly small enough to be able to make comparisons between landscapes without any need for statistical tests.

Impacts of applying the pesticide varied considerably between landscapes (Table 4). For 1X toxicity impacts varied between -0.027 to -0.143 for abundance, and -0.018 and -0.228 for occupancy. The range for 10X toxicity was -0.113 to -0.246 and -0.054 and -0.501 for abundance and occupancy respectively. Summed AOR scores provide an estimate of the total impact. Summed AOR for 10X toxicity was correlated with summed AOR for 1X, and was larger by a factor 2.14 (regression y = 2.14x - 0.0301, R^2^ = 0.5, n = 10). Impacts were highest in Mors, Næstved and Odder, and lowest in Tønder. The lower impacts seem to correlate with more extensive farming in this area.

**Table 4:** Occupancy, abundance and summed AO scores for each landscape after treatment with an endocrine disruptor set at 1X and 10X toxicity. Es = Esbjerg, Hi = Himmerland, Ka = Karup, Ko = Kolding, Lo = Lolland, Mo = Mors, Na = Næstved, Od = Odder, To = Toftlund, Tø = Tønder

| Endocrine Disruptor 1X Tox. | Es | Hi | Ka | Ko | Lo | Mo | Næ | Od | To | Tø |
| --- | --- | --- | --- | --- | --- | --- | --- | --- | --- | --- |
| Occupancy | -0.077 | -0.095 | -0.167 | -0.112 | -0.028 | -0.228 | -0.191 | -0.143 | -0.220 | -0.018 |
| Abundance | -0.070 | -0.070 | -0.030 | -0.086 | -0.143 | -0.033 | -0.032 | -0.056 | -0.027 | -0.049 |
| Summed AO | -0.147 | -0.165 | -0.197 | -0.197 | -0.170 | -0.261 | -0.223 | -0.199 | -0.247 | -0.067 |
| Endocrine Disruptor 10X Tox. |  |  |  |  |  |  |  |  |  |  |
| Occupancy | -0.221 | -0.233 | -0.179 | -0.343 | -0.161 | -0.481 | -0.501 | -0.424 | -0.370 | -0.054 |
| Abundance | -0.152 | -0.160 | -0.123 | -0.185 | -0.246 | -0.136 | -0.149 | -0.173 | -0.119 | -0.113 |
| Summed AO | -0.373 | -0.393 | -0.301 | -0.528 | -0.408 | -0.617 | -0.649 | -0.598 | -0.489 | -0.167 |

To attempt to make a simple model that could be used to predict impact, the sum of the AOR scores was used as a measure of impact and plotted against baseline hare density and landscape descriptors for both 1X and 10X toxicity (Fig. 2). In 1X toxicity scenarios density was the best determinant of impact (R^2^ = 0.52), followed by arable area (R^2^ = 0.49), pesticide treated area (R^2^ = 0.38) and lastly treated area multiplied by number of applications (R^2^ = 0.25). However, the pattern was quite different in the 10X toxicity scenarios. R^2^ for density was 0.03, arable area 0.35, treated area 0.88, and treated area multiplied by number of application 0.86.

**Figure 2.**
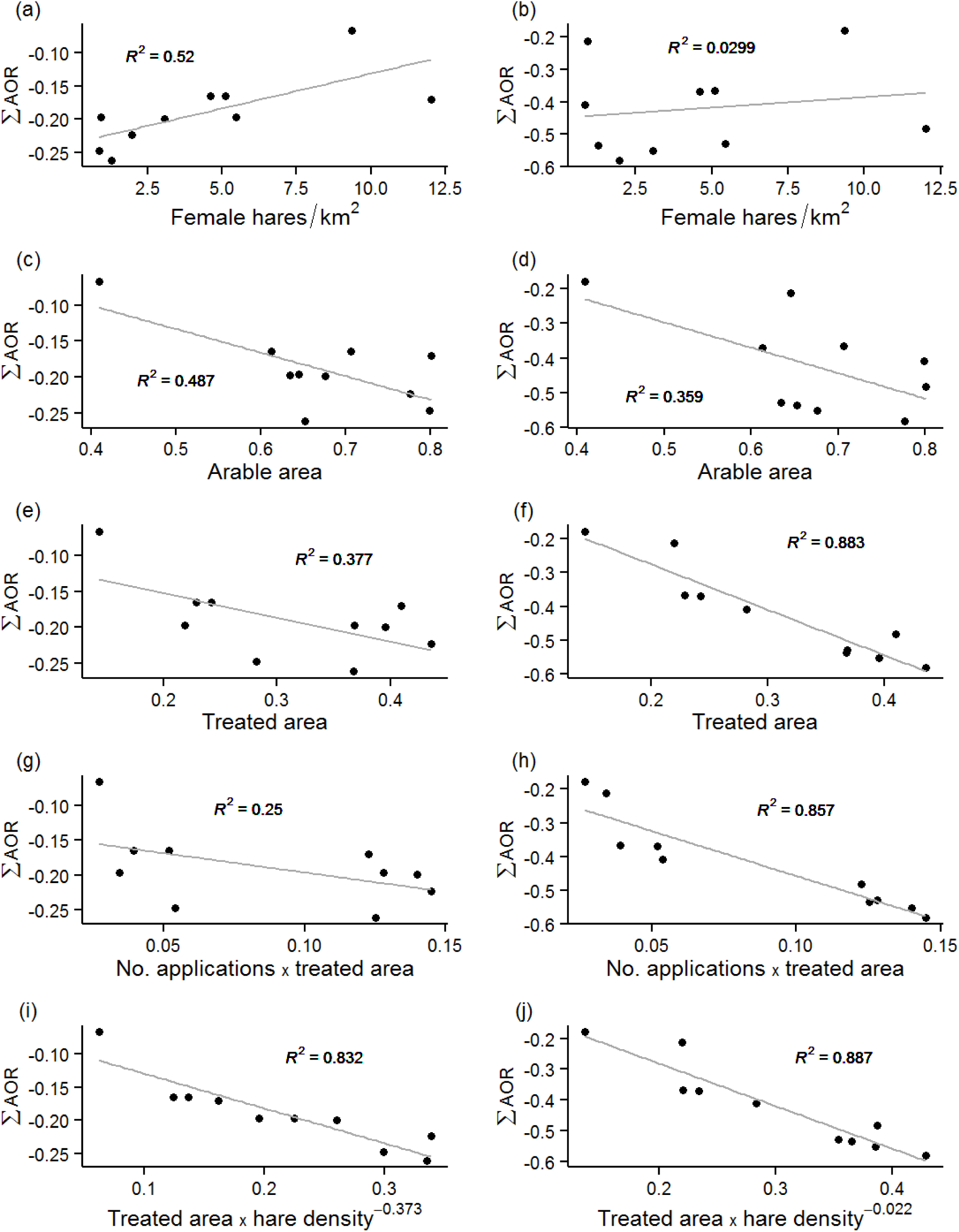
Summed AOR plotted against model data inputs for 1X (a,c,e,g,i) and 10X (b,d,f,h,j) toxicity scenarios and 10 landscapes with linear regression and associated R. a - b female hare density, c - d total arable area, e - f pesticide treated area, g - h pesticide treated area multiplied by landscape specific mean number of applications, I - j best fit model combining hare density and area treated.

In both cases density was combined with the best correlated landscape descriptor to try to improve model fit using linear, geometric and exponential combinations. Here our aim was to find the best way to combine these parameters to create a good meta-model for the ALMaSS simulations. The resulting best overall model was:

Summed AOR *as* = *a*℮*^d^i^,* where *a* is the proportion of area treated, and *d* is the density (adult female km^−2^) and *i* is a constant.

R^2^ (n=10) for the fitted model was high (0.83 & 0.89 for 1X & 10X toxicity), thus impact could be predicted using the regression equation of this model. This, however, would require both treated farm area and hare density as inputs, and the equation for the model was rather different in 1X & 10X versions, with *i* having values of 0.373 and 0.022 respectively. However, the fit for 10X toxicity was equally good based on treated area only (Fig. 2).

To further analyse the extent landscape structure and farming influenced the impact of the pesticide the 10X simulation was run for all possible combination of 5 landscapes and their farming. The resulting hare densities varied slightly more than the original 10 landscapes from virtual extinction in all Lolland landscape runs (except when using Lolland farming), to 14 adult females km^−2^ for all other landscapes with Lolland farming. There was a clear separation between both farming effects and landscape effects on the hare densities (Fig. 3). Further analysis was performed using AOR scores for all landscape combinations with pesticide compared to the same combination without pesticide.

**Figure 3:**
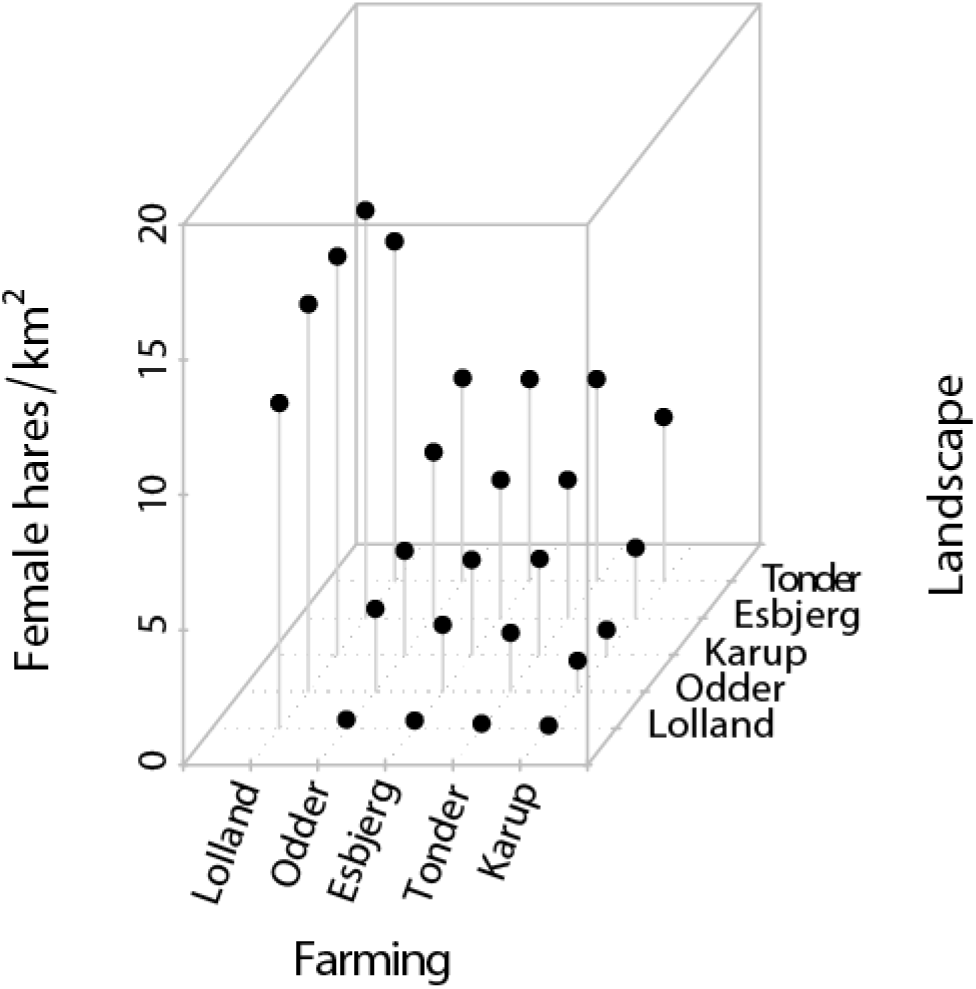
Adult female hare densities plotted for all 25 farming and landscape combinations run. Lolland landscape has very low hare numbers unless farmed using real Lolland farming, which his highly hare beneficial in all landscapes.

Using the same model as used to predict summed AOR scores for the 10 baseline landscapes did not predict impacts as well for this larger data set (R^2^ = 0.42, n= 21). The four low density Lolland landscapes could not be fitted using the model chosen and were ignored as outliers. There was no clear pattern discernible from the fit except an indication that the Tønder landscape deviated more from the linear fit than the others. It was not easy therefore to predict impacts with accuracy based on a knowledge of area of pesticide use and hare density, even though in the smaller landscape set this seemed promising.

AOR plots for scenarios where landscape/farming combinations were tested showed that when considering impacts of a pesticide on a population that both landscape and farming can have very large influence on the ERA endpoints. In the Tønder landscape the impacts of pesticides were low for all farming regimes tested, with impacts primarily on abundance of 0-5% (Fig. 4). In contrast, impacts on the Lolland landscape were in general very large reductions in occupancy due to the low population size resulting from 4 out of 5 of the farming regimes, and large impact on abundance in the one farming regime that gave high hare densities. Farming regimes had different influences in all the different landscapes, although the impacts in landscapes with Lolland farming were all primarily in reduction of abundance (as a result of the high overall population density). Overall, with the exception of the Tønder landscape, the results of the ERA were not constant in any of the landscapes or farming regimes. These variations in impact were large enough to result in a number of these landscape/farming combinations to fall below the arbitrary -0.1 summed AOR threshold assumed in the initial real landscape/farming combination scenarios. These combinations all had either Tønder landscape or Lolland farming (Fig. 4).

**Figure 4.**
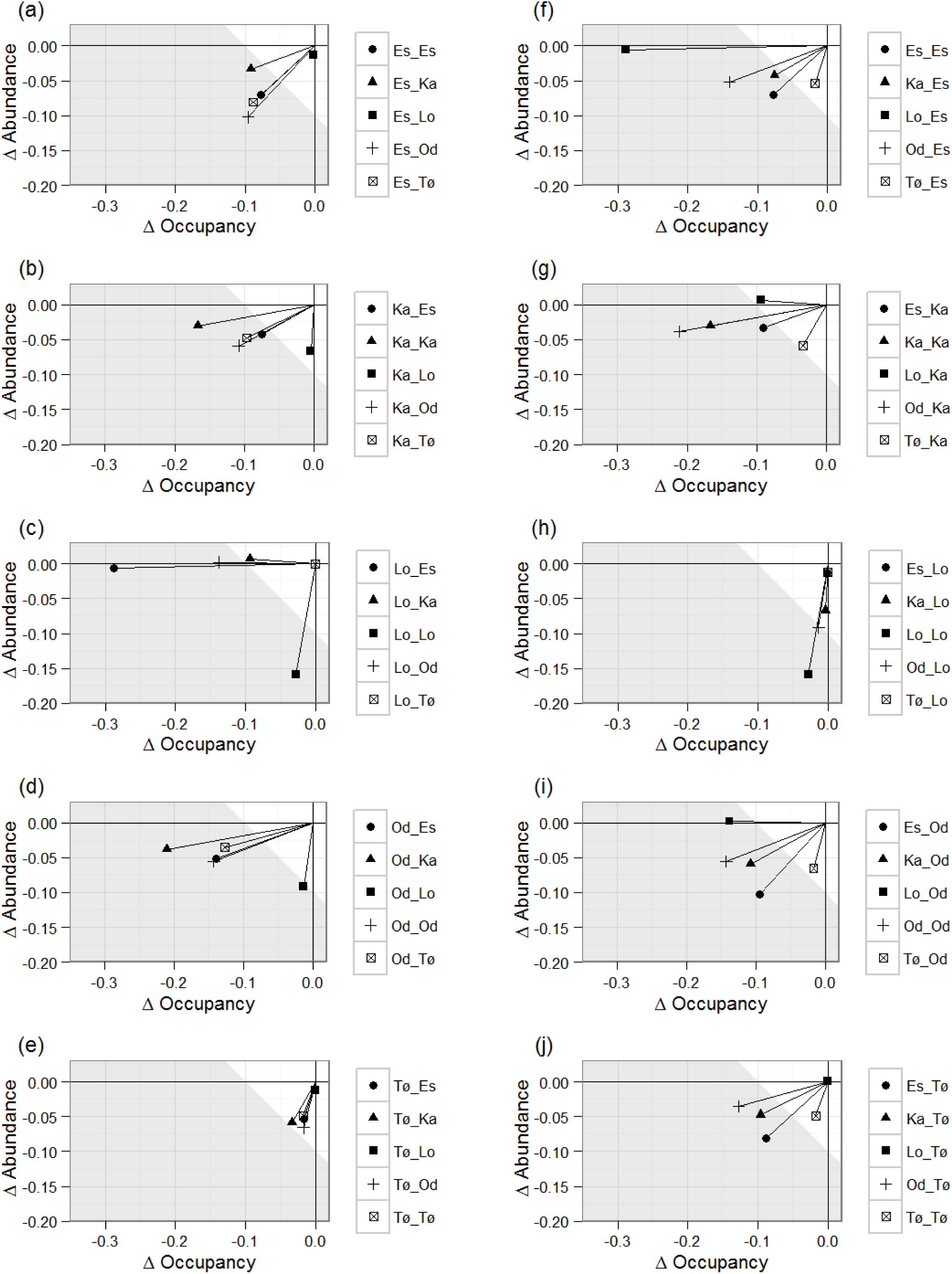
AOR plots for each landscape/farming combination tested. Each pair of plots shows the variation resulting from either using different farming on a single landscape (landscape) or applying the same farming to different landscapes (farming). a – Esbjerg landscape, f – Esbjerg farming, b – Karup landscape, g – Karup farming, c – Lolland landscape, h – Lolland farming, d – Odder landscape, i – Odder farming, e – Tønder landscape, j – Tønder farming. The white area exemplifies a specific protection goal of no more than -10% change in the sum of occupancy or abundance changes.

## 4. DISCUSSION

The landscape generation method provided a chance to evaluate the extent to which a mammalian pesticide ERA was affected by the type of landscape and the farming regime in force on that landscape. Generation of 10 × 10 km landscapes takes only a few hours of computer time on standard PC architecture (2.8 GHz Intel Core-i7, 32GB), and is almost totally automated once rules for landscape generation have been decided upon. Using this methodology allowed creation of 10 different and detailed examples of Danish agricultural landscapes together with their respective farming regimes. This provides the basis for generating scenario input and in conjunction with suitable measurement endpoints and population models creates a relatively easy method for creating landscape-specific ERAs.

Impacts in ERA are likely to be needed to be measured in both changes in abundance and also distribution (EFSA Panel on Plant Protection Products and their Residues (PPR) 2015). It is therefore important that both are incorporated in the measurement endpoints. This can be achieved using the AOR–scores (Topping, Craig et al. 2015). The advantage of this approach is that it is very easy to make comparisons relative to a baseline condition and to assign a threshold for unacceptable effects. Here we have assumed a threshold of -10% change, and that abundance and occupancy have equal weight (e.g. Fig. 4). The use of summed AOR scores as a measurement endpoint provides a way to combine both changes in range and abundance into a single metric. Summed AOR scores provide the easiest way to work, since they can be directly compared to the trigger value set by the SPG. However, summing the scores makes some assumptions; chiefly that equal weight is assumed to be given to changes in both. This assumption can be violated easily if the grid size used for occupancy is unsuitable (too large and there is no sensitivity, too small and abundance and occupancy are subject to stochasticity). It may also be violated if there is some *a priori* ecological or social reason that distribution is more or less important than density. If the latter is the case, then it would be possible to weight the summed AOR scores accordingly before assessment.

When carrying out the ERA for the fictitious endocrine disrupting pesticide using summed AOR scores there was a general agreement between landscapes that the 10X higher toxicity scenario had higher impacts than the 1X toxicity scenario. However, impacts were approximately doubled, nowhere near the 10X factor on toxicity. Similar responses have been seen before (Dalkvist, Topping et al. 2009), where vole population ERA was influenced at least as much by ecological factors as by toxicological ones. This should not be a surprise. Three main factors will dampen responses when considering a population approach. The first is that there may be local buffering of impact as a result of local density-dependent processes resulting in faster reproduction at lower population levels (e.g. Kramarz, Banks et al. 2007). The second is that the impacts are spatial, and mixing of the population is not complete, hence impacts can be reduced if recolonization to continually treated areas is slow. Finally, a threshold response effect (e.g. preventing reproduction) means that, once triggered, further increase in toxicity has no effect. Of these three, the latter is likely to be the most important in this case, since hares will generally move quickly into unoccupied suitable areas and become exposed. In this case the population responses will also be delayed since hares are long-lived if they reach adulthood (Abilgaard, Andersen et al. 1972), and temporarily reduced reproduction may be compensated for during the lifetime of the hare.

Impact varied tremendously between landscapes and farming. This was a consequence both of the pattern of exposure and of internal population processes. The effect of internal population processes can be clearly seen in populations exposed under the Lolland farming regime where high population levels buffered changes in occupancy, leading only to impacts on abundance. However, exposure must also differ between landscapes since the populations were of approximately the same size, but magnitude of impacts differed between landscapes with Lolland farming. Area of treatment was not a good predictor and in some cases was barely correlated with overall impact. It is important to be aware, however, that these ecological drivers are probably more important than precise toxicity and exposure measures in cases where effects are certain and the implication of these effects at the population level are the main point of interest. In cases where prediction of individual toxic effects is in focus, then toxicity will have a major influence on impact and toxico-kinetics and toxico-dynamics will play a more important role. This would be the case if the pesticide caused acute rather than chronic effects and individual deaths were unacceptable; for example, in mammal and bird risk assessment where no individual mortality is used as a surrogate protection goal for no visible mortality (EFSA 2009).

The variation linked to both farming and landscape makes matching the correct farming with the correct landscape critical to prediction of impact. Some mixed landscape/farming settings altered impacts significantly and would in an ERA lead to errors in estimating risk. Similar hare sensitivities to management were also seen for specific management changes as well as farming regime changes between organic and conventional management forms (Topping 2011). From a regulatory ERA perspective this raises a complex issue, i.e. which landscape/farming regime should be used to base the assessment upon. A basic principle of regulatory ERA is that scenarios are based on a reasonable worst-case scenario (e.g. EFSA Panel on Plant Protection Products and their Residues (PPR) 2014), but how should these worst-case scenarios be identified? Both farming and hare density were the major determinants of the impact, but prediction of these requires knowledge of the landscape structure, and density relies on farming. It is quite possible to make an erroneous judgement here based on observation from the real world. For example the Lolland landscape had the highest predicted hare population, and also has the highest agricultural hare densities in Denmark, thus it probably would not be considered worst case. However, unless the specific farming regime of Lolland is used in the scenario, populations there were predicted to be the lowest in any simulations. This context dependency is unfortunate, but is a feature of ecological systems (Noda 2004, Chamberlain, Bronstein et al. 2014, Paterson, Dick et al. 2015). Reasonable worst-case is therefore not easy to predict. Accurate assessment is also clearly a case of ensuring the correct landscape and farming is used, since simplifications and generalisations may have unexpected consequences.

Context dependency is also very clearly shown in the meta-models used to analyse correlations between summed AOR and scenario descriptors. Creation of a meta-model is clearly a good idea given the complexity of developing and running the scenarios, since prediction of impacts would be much simpler. However, this example demonstrates the danger in extrapolating results from one (even replicated) scenario to another. In this case only toxicity was changed, but this dramatically altered the overall model and predictions based on one toxicity failed to model the other accurately. This result may indicate that the practice of meta-modelling (also called Emulation) of complex agent-based simulations may need to be applied with caution to scenarios that are not represented by the original training dataset. These approaches are typically currently used to evaluate sensitivities of model (e.g. Happe, Kellermann et al. 2006, Parry, Topping et al. 2013), but the idea of using a simplified meta-model to predict, in this case population, impacts under new conditions would be attractive, and clearly could be misleading.

This study uses a single species of mammal and a single pesticide impact on litter size. Whilst the results clearly demonstrate context dependency on landscape and farming, and therefore should be of concern in regulatory ERA, the extent to which these results can be extrapolated to other species and effects is currently unknown. The critical interactions which determined the landscape and farming influence on the ERA here were those determining hare density at low (1X) application rates, but changed to area treated at 10X toxicity. This is probably a fairly general pattern. Medium and low effects will be buffered by a highly fecund population, and large effects by spatially distributed sub-populations, at least in highly mobile species. However, the resources required to produce these highly fecund/high-density or spatially distributed populations will vary with species. In the hare, large areas of low intensity grazing provided a solid buffer in the Tønder landscape, whereas the high-density populations occurred in Lolland due to type of farming. Similarly, the precise impact (e.g. reproductive effect, mortality, behavioural changes) will interact with the population dynamics to change density and distribution in different ways in different species. Therefore, whilst the conclusions indicating the importance of context dependency on ERA are likely to be general, the specifics will be species and stressor-effect dependent. Developing a large number of species models and carrying out a widely varying set of ERA scenarios may provide general rules to determine for which species context is critical and for which generalisations can be made but was beyond the scope of this study. However, some preliminary tests (not presented) using a toxic standard test of direct mortality as suggested by Schmitt, Auteri et al. (2015), indicated that similar interactions with landscape and farming might be expected with other modes of action.

Context dependency is a problem for regulatory ERA where the ideal approach would be a set of standardized scenarios, for instance as is done for surface water in Europe (FOCUS 2001). A single scenario, for in this case Denmark, would only work at a very large scale where all possible conditions would be encountered. However, this would not be feasible from a simulation perspective, and would also run the real risk of being unable to detect impacts at lower spatial scales that might be a cause for concern (e.g. if regional populations became extinct). One alternative based on the approach used here would be to select a range of representative areas and use these as a basis for screening for unacceptable inputs. This would have been very difficult previously; however, the methods used here for extracting the information necessary for ERA at this scale are now available and can work with EU subsidy scheme information and GIS layers. This provides a very important tool for landscape scale risk assessment in the future, since previously developing landscape data of this resolution as well as linkage to farming was very difficult and time consuming; previous versions of ALMaSS relied on hand-digitised maps taking two months to produce. Future improvements to this process would accrue from access to the pesticide use data collected under the Sustainable Use Directive (EU Directive 2009/128/EC), which requires farmers to record their pesticide usage.

## 5. CONCLUSIONS

The need for landscape scale risk assessment is becoming more widely accepted. However, tools to achieve this are scarce. Models are being developed for this scale of assessment, but until now there was no quick and reliable way to generate landscapes detailed enough to run these simulations on without recourse to laborious procedures. The methods for landscape creation presented here make creation of landscapes for risk assessment feasible based on EU subsidy information and planning maps.

Both landscape and agricultural regime influence the result of an ERA for and endocrine disruptor on populations of hares considerably. Area treated was a poor indicator of impact. The major determinant of the impact was the baseline population level, but this is difficult to predict without accurate simulation models, and changes in farming can have very large impacts on this metric. Accurate landscape and farming information is therefore necessary to determine impacts of pesticides on hare populations. The extent to which similar effects can be expected with other species and models of action is unknown, but this seems likely.

Since prediction of a reasonable worst case scenario is difficult from landscape or population metrics measured in the real world, future ERA may need to make use of multiple landscapes (or “scenarios”) as a screening tool to avoid locally unacceptable risks. Given the advances in landscape generation this approach could now be feasible Europe wide. However, the results caution against the use of meta-modelling to represent new landscape/farming/species combinations which could give spurious results.

## 6. ACKNOWLEDGEMENTS

We acknowledge the support of the Danish Environmental Agency research grant for the project “Udvikling af et værktøj til risikovurdering af pesticider, der tager højde for rumlige og tidsmæssige processer, som eksisterende metoder ikke kan håndtere” which supported the development of the landscape generation methods. We would also like to thank Steen Gyldenkærne for helping with access to the GLR and CHR databases and Michael Stjernholm for GIS support.

## Appendix 1 Generation of model landscapes for ALMaSS

The process of generating a complete simulation landscape for ALMaSS is divided into two main tasks:

1. Farm classification (to classify all farms in Denmark into a number of general farm types)
2. Generation of landscapes as input for ALMaSS

### Farm classification

A program was written in C++ to classify all farms in Denmark into general farm types (http://www.ecosol.dk/MSTProject/Documentation/FarmClassification/index.html). The program classifies all farms based on a combination of the crops they are growing using data obtained from the General Farm Register (“Det Generelle Landbrugsregister” - GLR), and on the animals they have which is data from the Central Livestock Register (“Det Centrale HusdyrbrugsRegister” - CHR). The GLR is a compilation of the data submitted by the farmers in support of EU subsidy payments. The CHR is a register of all agricultural animals maintained primarily for purposes of disease control.

By combining crop and animal information it was possible to identify major farm types such as pig, arable, or dairy farms. Some less common types are also identifiable e.g. farmers that grow sugar beet on contract. In addition to this information the GLR also indicates whether a farm is organic or not and the overall farm size. This extra information provides the basis for the classification. Rules used to classify the farms were needed to be very general because real farms tend not to fit neatly into pure farm type rules (e.g. many arable farms have grazing because they have some animals e.g. horses or a few animals for their own consumption. The rules used were:

1. Farms with large proportion of vegetables (minimum 0.5) and larger than 2-ha were organic or conventional Vegetable farms, otherwise if small were classified as ‘other’.
2. Farms with a proportion of potatoes or sugar beet not less than 20% were Potato or Beet farms respectively.
3. Farms with animal (cows, sheep and pigs) transformed to standard animal units that have fewer than 20 animal units and an area less than 20 ha were designated as Hobby farms (<25 ha is typically part-time or hobby (Levin 2006), so 20 ha will reduce the chance of misclassifying commercial farms).
4. Farms with animal units above 20 and cattle + sheep above 75% they were designated as Cattle farms
5. Farms with animal units above 20 and pigs above 75%, or crop area of grazing pigs above 15% were designated as Pig farms.
6. Farms with animal units above 20, but not pig or cattle farms, were designated as Mixed Stock.
7. Farms with no animals registered but with large areas of grazing were assumed to be either Cattle farms or Mixed Stock farms depending on whether grazing area was above 40% or 20-40% respectively.
8. All remaining farms must have been Arable farms (i.e. large area with few or no animals and little or no grazing).
9. All farms except ‘other’ could be designated as organic or not dependent upon the information on that farm in the GLR giving a total of 17 farm types possible.

For each farm type, the mean proportion of the farm crop area was calculated for each crop. Crops with less than 1% share of the area of a farm type were ignored and the rest used to create a farm rotation for that classification. It was assumed that the rotation could be represented by 100 crops (1 crop for each 1%). The order of crops followed typical agronomic practices and issues such as late harvest leading to impossible sowing conditions were controlled by the built in ALMaSS farm code (see Appendix 2). The result is a pattern of changing crops on a field that matches the overall crop distribution pattern for that farm type precisely over 100 seasons. Viewed on a larger scale crop distributions will therefore be overall correct at any point in time, although the actual crop grown on a single field will not replicate reality. This method does not, however, take into account differing soil types between fields, which in reality would restrict some crop distributions.

### Generation of ALMaSS simulation landscapes

The aim of this task is to generate a land cover raster map with complete coverage; hence all cells must be classified in accordance with their landscape element type. In most cases existing land cover maps are in a coarse spatial resolution (e.g. 100m * 100m; Corine Land cover 2006, EEA 2013) and landscape elements are often broadly categorized (e.g. ESA 2014). Thus for application in individual based modelling, such maps are inadequate. Alternatively manual digitization from areal images can produce maps of sufficient resolution and detail, but this is extremely time-consuming and therefore often not feasible for larger areas (i.e. several square kilometres). However, in many cases highly detailed vector maps are used for e.g. landscape planning and nature conservation purposes and these maps are increasingly becoming publicly available (Europoean Union Open Data Portal – open-data.europa.eu/en/data). Here we make use of a large number of such vector maps and combine them into a single raster map with high spatial resolution as well as with a large number of landscape elements. However, using a large number of different data sources can result in inconsistencies if maps have been made independently and/or if they have been made in different points in time. In most cases these inconsistencies will relate to the spatial alignment of vector layers resulting in either overlaps or gaps between features that are actually adjacent to each other (Figure 1).

**Figure 1.**
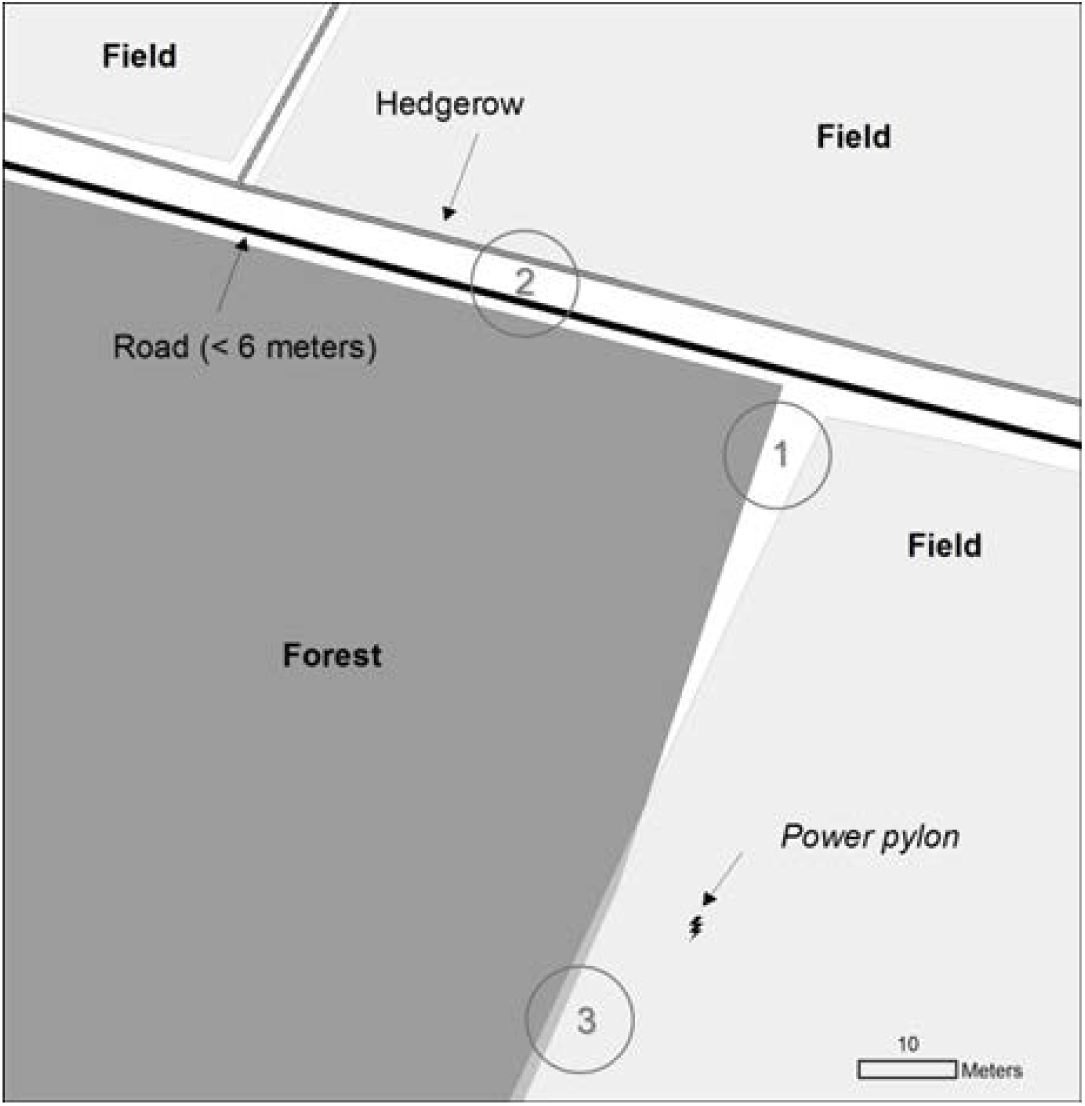
Examples of the three basic vector data types (polygons, lines and points) used as input to the land cover map. Vector data have been digitized for different purposes, in different points in time and they vary in spatial accuracy and detail and in the way the geometry is represented. The figure shows some of the common problems encountered when converting a number of vector themes to a surface covering map. (1) Gaps between polygons. (2) Lack of dimension, i.e. points are per definition dimensionless and lines have length, but not a width. This is obviously a cartographic abstraction and a land cover map need to address the exact extent of a feature (e.g., the width of a road, the actually area covered by a solitary tree etc.). (3) Spatial overlap. Vector layers differ in spatial accuracy which results in gaps or in overlap.

Additionally certain vector types, such as points and lines are dimensionless and therefore decisions about their dimensions needs to be made in order to obtain a meaningful mapping of these in a raster format (Figure 2). When working at high spatial resolution, these issues can be quite substantial and needs to be dealt with in order to obtain a surface covering land cover map. In the remainder of this section we describe methodologies to process a large number of input vector data to produce a surface covering land cover map while taking the abovementioned problems into account.

**Figure 2.**
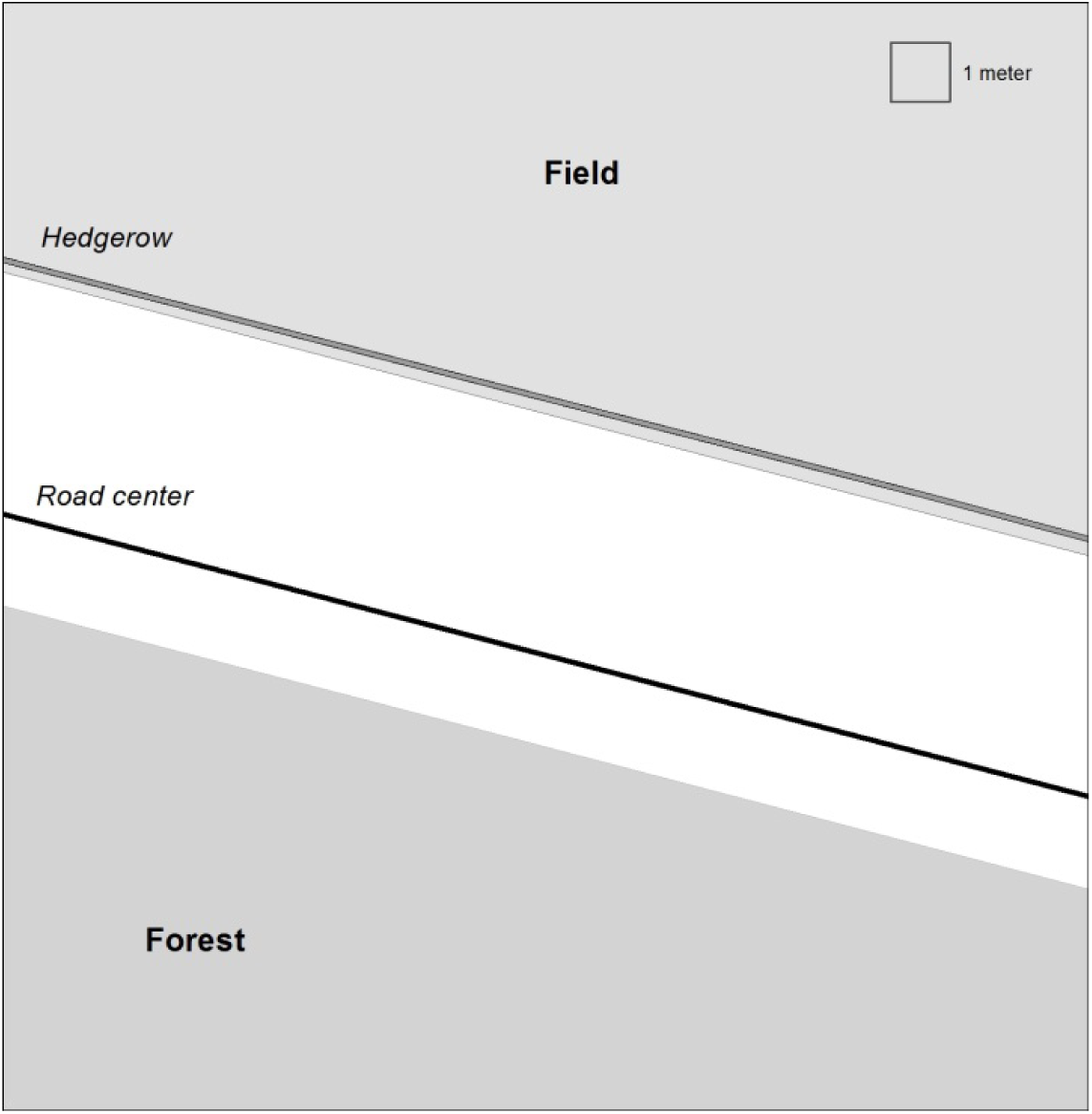
Part of figure 1 at finer scale showing gaps between polygons and lack of dimension. A forest- and a field-polygon with empty space in between, but two line features (a road and a hedgerow) will supposedly fill this gap. In this particular case, it is known that the road is a medium sized with a width between 3 and 6 meters. There is no further information about the hedgerow. The map reveals some problems: The middle of the road is not centred in the gap between the forest and the field and the centre of the hedgerow is situated inside the field polygon and divides it in two.

The overall process to generate the land cover map follows three steps: 1) Convert the input vector data to raster format, 2) combine individual raster layers into thematic maps (e.g., all road types, paths and railway tracks in a transportation theme), 3) stack these thematic maps in a reasonable order (i.e. roads on top of fields etc.). Below we first describe the input data, and then the three steps involved in generating the final map are described in detail.

#### Input data

We used publicly available vector map layers from the Danish common public geographical administration data (GeoDanmark data, downloaded 2012, http://download.kortforsyningen.dk). The vector layers were used to map 41 different landscape features (see table 1). The Danish AgriFish Agency (DAFA, under the Ministry of Food, Agriculture and Fisheries) provided vector maps of individual agricultural fields. Lastly, where a pixel was not covered by a field or the GeoDanmark layers (approximately 3-5% of the area), we used the Area Information System (AIS) data, which is a surface covering map with 45 different land cover types for Denmark (Stjernholm et al., 2000). The AIS map is based on data and satellite imagery from the late nineties, but is, nevertheless, the best surface covering map available.

**Table 1.**
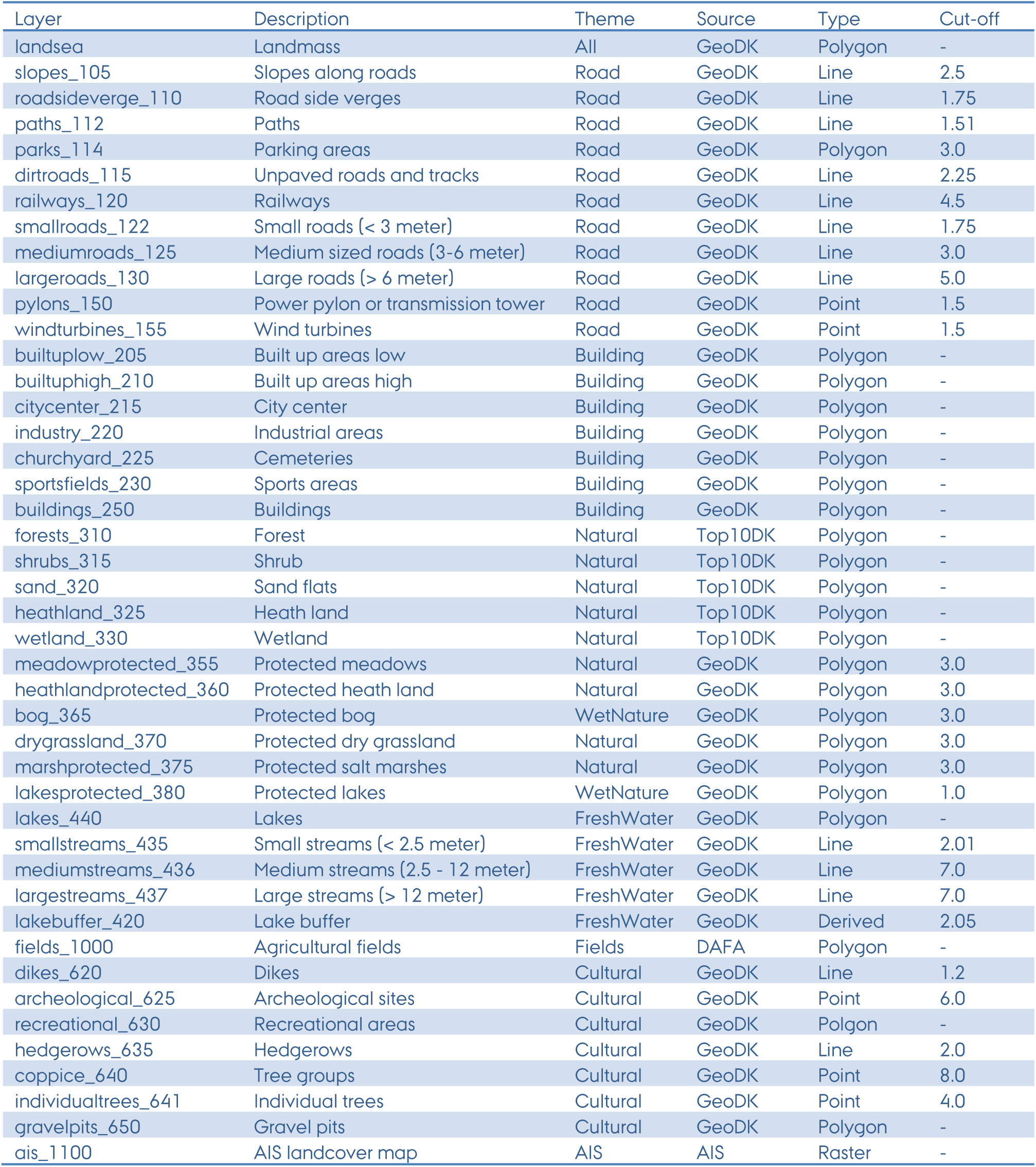
Description of the individual layers used in the final map, the theme in which they are grouped, their data sources and the original data type. Layer names follow the convention shortdescription_numericvalue, e.g. the layer slopes_105 is the layer with slopes along roads and 105 is the numeric value indicating presence of the feature.

The Danish AgriFish Agency (under the Ministry of Food, Agriculture and Fisheries) maintains a map of all fields and a database of crops grown in Denmark. The database is updated annually. Farmers are obliged to report for each individual field the crop they intend to grow the following year. The data set used for this study stems from 2013 where more than 45.000 farmers contributed to the database. The data set makes it possible to identify the owner (or manager) of each field and the actual crop grown on it.

The ALMaSS landscape simulator modifies the actual production in each field based on the dominant soil type. The Danish centre for food and agriculture (DCA) at Aarhus University maintains soil classification maps and we used the 1:200.000 map for this study (downloaded from http://dca.au.dk/forskning/den-danske-jordklassificering/). The soil classification map was rasterized and field polygons were overlaid to determine the dominant soil type for each field.

#### Making the map

All handling and analysis of spatial data was done using Python 2.7 and the python library arcpy to access ArcGIS features (ESRI 2010). For documentation of each individual arcpy tool used see help.arcgis.com (search for: ‘What is ArcPy?’). The entire process of producing surface covering land cover map has been programmed in a python script that is freely available on Github (https://github.com/flemmingskov/python-landscapegen/tree/PrepForPub). In addition to the data described above an outline of the simulation area is needed before running the python script. The outline needs to be rectangular, in raster format and have the desired spatial resolution (usually 1m by 1m). The outline will be used as a clipping mask to clip any of the data layers that extend beyond the simulation area (Figure 9).

##### The Python script

The python script is divided into 5 sections of which the first four makes the land cover map. The fifth is only needed if the map is prepared in order to run ALMaSS simulations. Each main operation is described here with references to figures where relevant.

In the first section (Setup) the libraries needed are imported, paths to input data and outputs are defined, the processing environment is defined and generation of individual themes and layers can be switched on or off. The script assumes a geodatabase to store output (Figure 9, top right).

The second section (Conversion), which constitutes the majority of the script deals with the conversion of the original vector data into raster format. For linear and point features the conversion process involves two steps. First calculate a raster with the Euclidian distance from the features (Figure 3) and second define a raster with the numeric value for the feature at a defined distance from the original vector feature and 0 beyond this distance (Figure 4; see Table 1 for detail about distance used for each of the layers). The numeric value chosen to indicate presence of the feature determines the hierarchy when later combining individual layers into themes. Thus at this stage care must be taken to ensure that numeric values within each theme are sensible. Polygon features are converted directly without adding a buffer, except for a few cases where the original mapping was inaccurate (mostly because the features mapped were difficult to delineate, e.g. the border between swamp and marshland).

**Figure 3.**
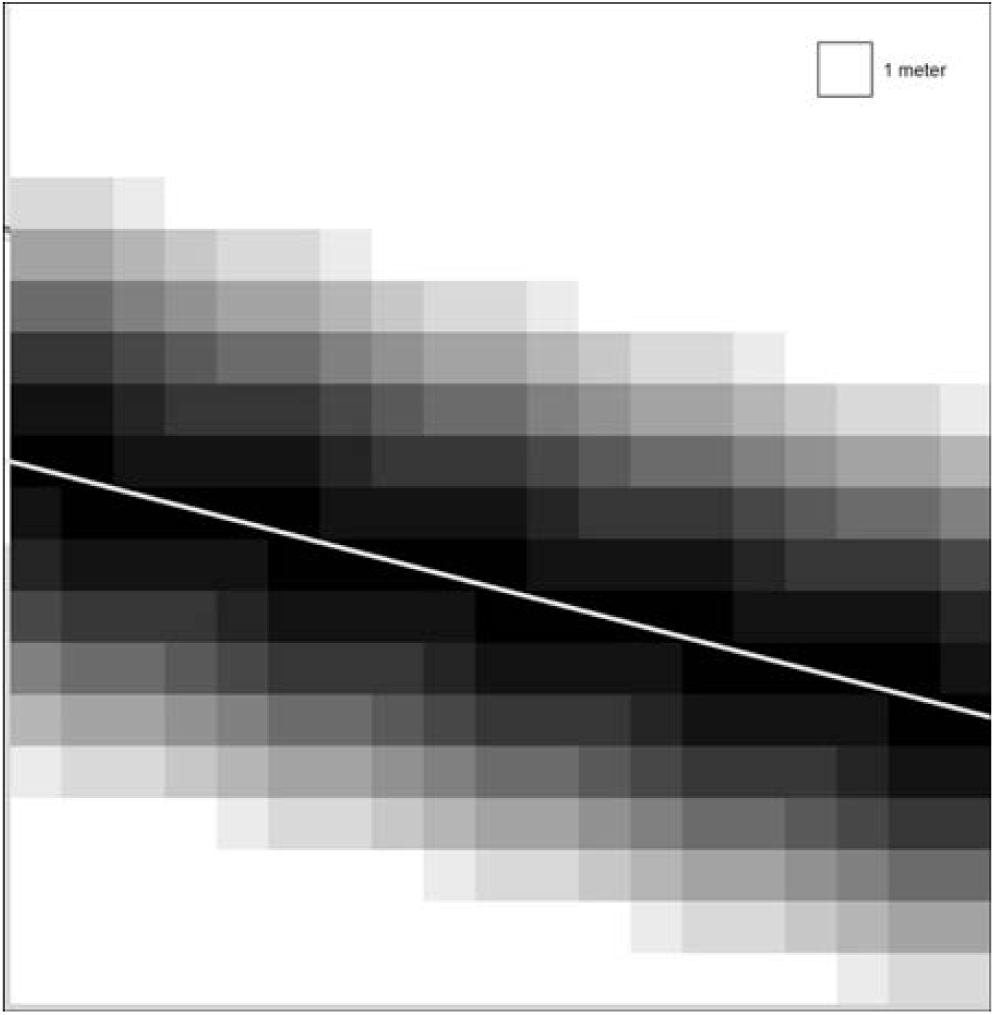
Conversion of a vector feature (here a line) to a raster layer. For each cell in the raster grid the Euclidean distance to the nearest point of the line is calculated (the darker the shading, the closer the cell is to the line). The next step is to choose a cut-off value to select the cells that will be coded as ‘road’ in the final land cover map.

**Figure 4.**
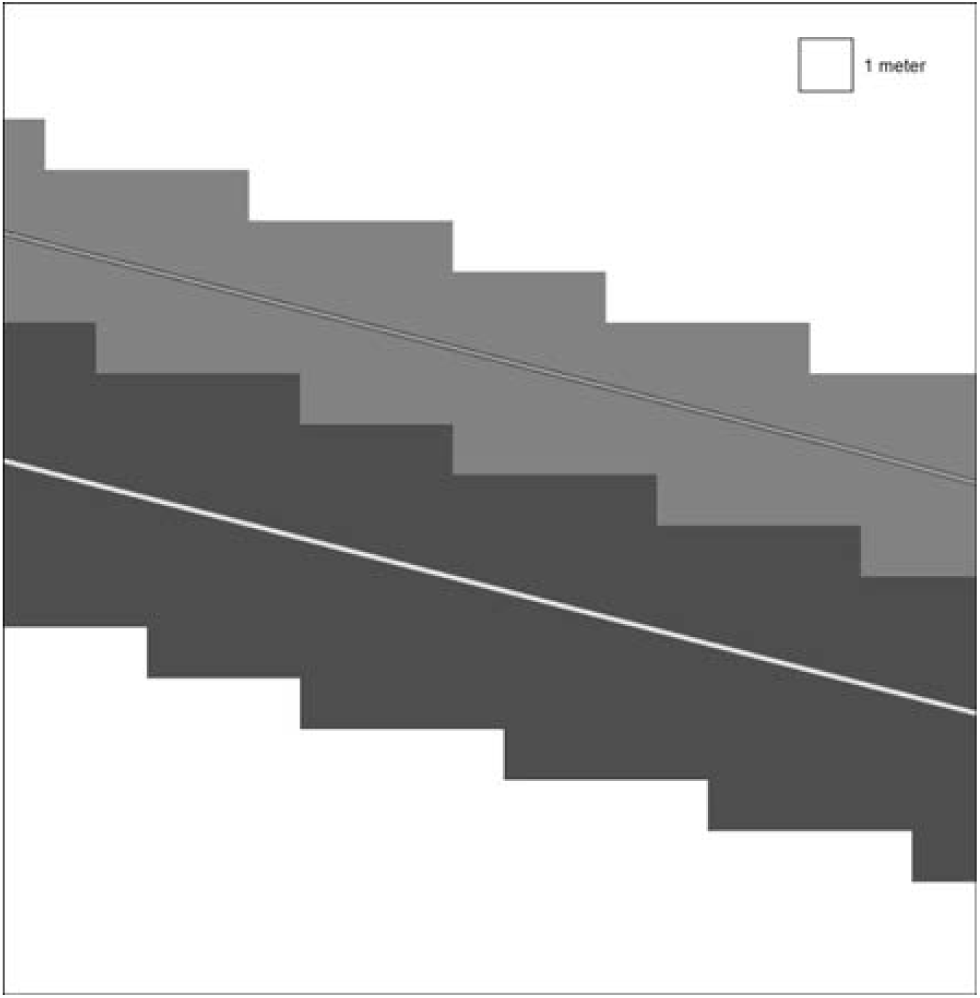
The final representation of ‘road’ and ‘hedgerow’ in the land cover map is shown here. The actual choice of a width for a line feature depends on the information available and the purpose of the map. In the present case it is known that the road is medium sized (between 3 and 6 meters); the highest of the two values was chosen and all cells <= 3 meters from the road centre was included as road. No information about the width of the hedgerow is available. Here an average width of 3 meters was chosen.

The third section (Themes) collects the raster layers into thematic maps (e.g., all road types, paths and railway tracks in a transportation theme etc., see Figure 5). In cases where two or more of the layers in a theme overlap, the layer with the highest numeric value is prioritized. For example if a large road (numeric value 130) intersects a small road (numeric value 122) the large road gains predominance and is shown on the final map (Figure 6).

**Figure 5.**
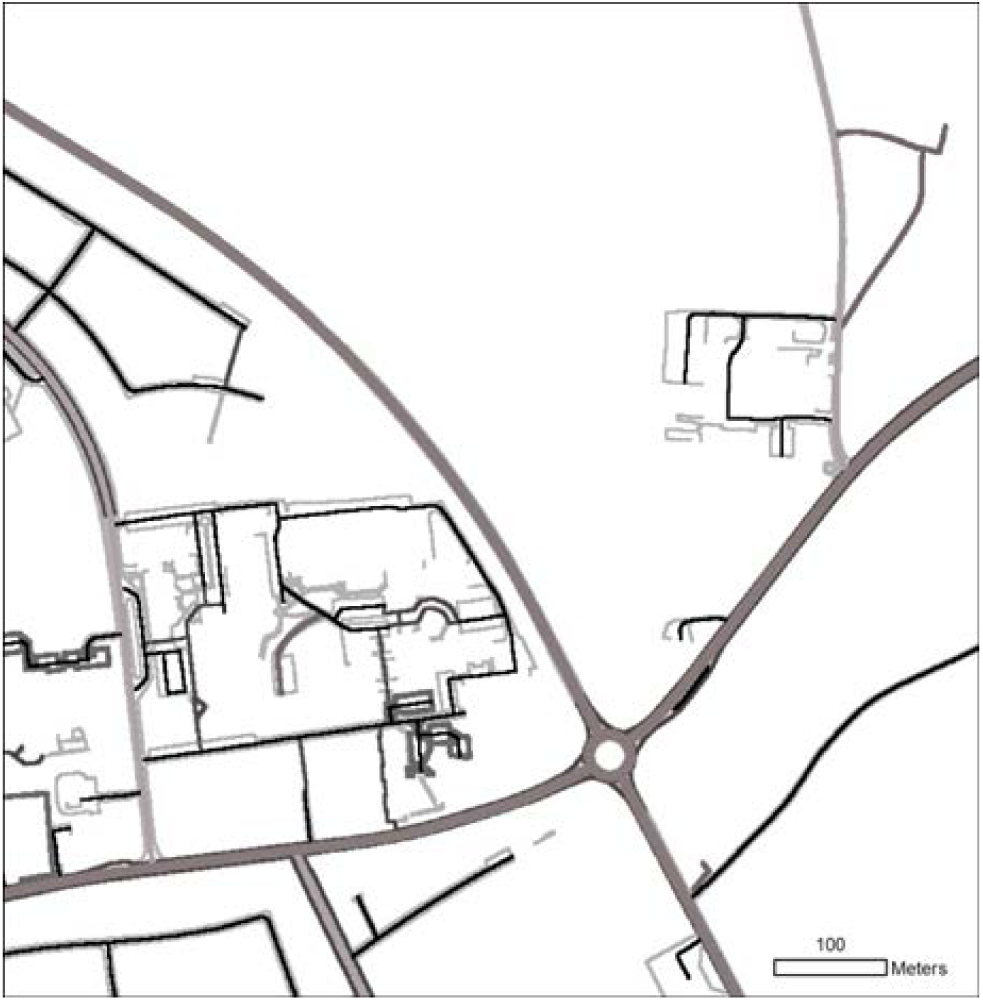
Example of a thematic map. The road network theme is made up of 11 individual raster layers. Each of these raster layers has a numeric code > 0 assigned to cells occupied by the feature in question (and 0 if the feature is absent) (see Table S2). The final theme is made by comparing all raster maps cell by cell and choosing the maximum value. Thus the numeric value rank features and determines which layers will occupy a given cell when more features are present. In the road network theme, for example, small roads (code 122) will precede railway tracks (code 120). The choice of codes is therefore important and depends on the purpose of the final land cover map.

**Figure 6.**
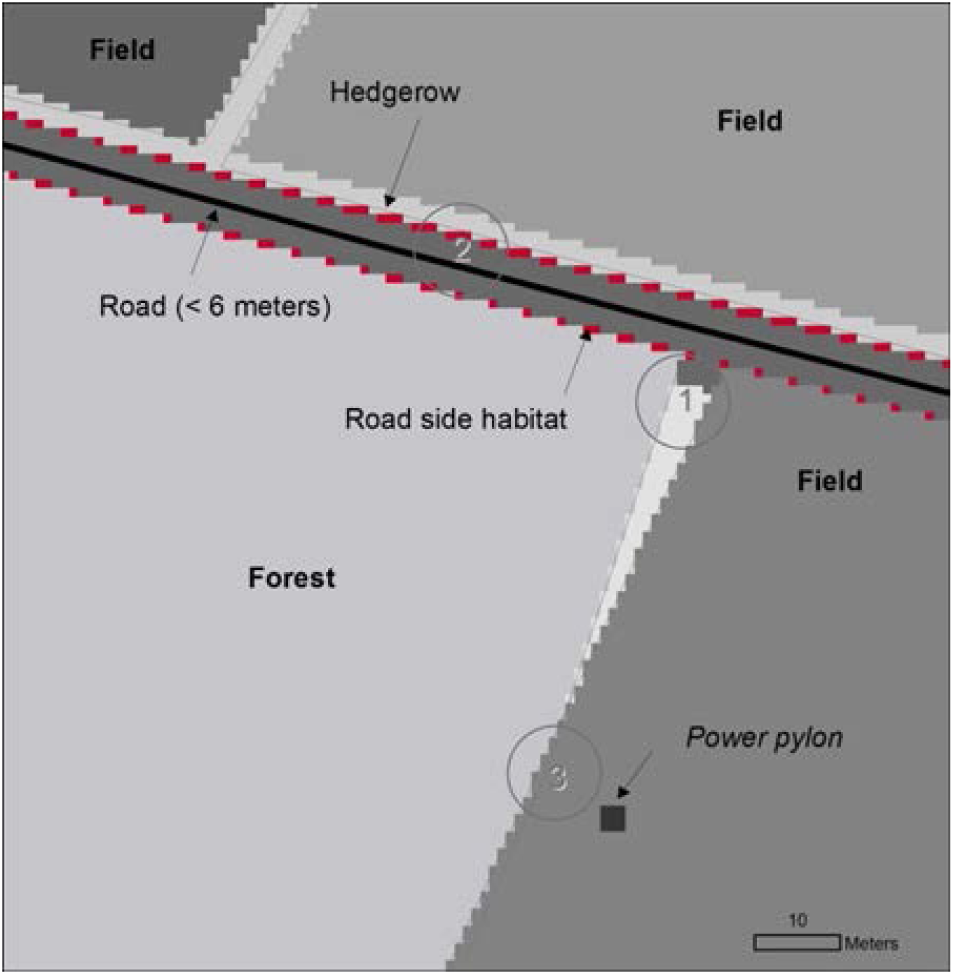
Map assembly and final mosaic map. The individual themes are put together according to a set of rules determining the order of stacking. The order depends on the purpose of the final land cover map and may be changed accordingly. The numbers refer to the problem areas described in Figure 1. The procedure is shown in Figure 7.

The fourth section (Stack) stacks the thematic maps in sequence such that the final map shows the ecological meaningful layers on top. E.g. the fresh water theme has to be stacked on to the fields to avoid artificial overlaps (Figure 7).

**Figure 7.**
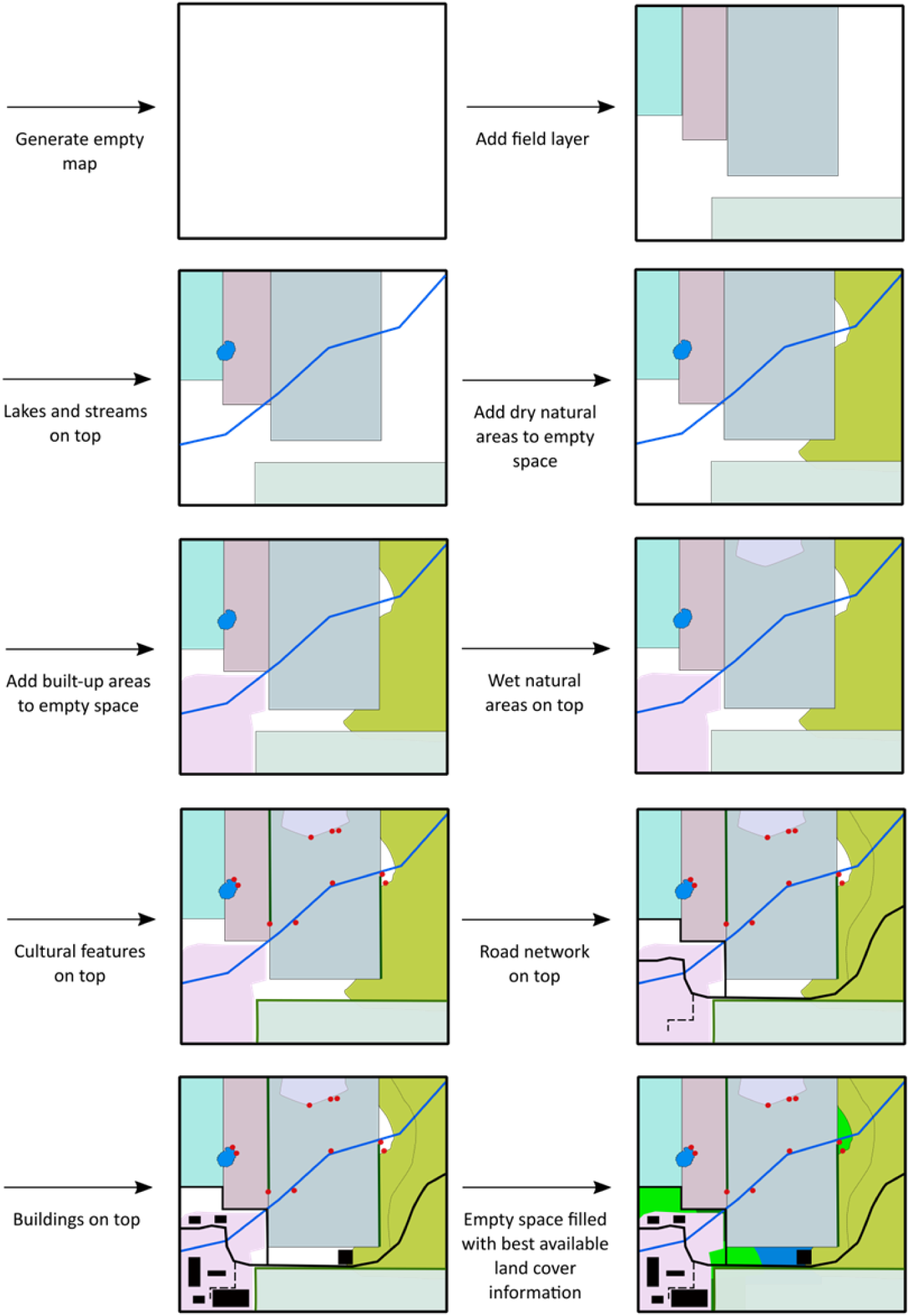
Stacking. Illustration of the stacking procedure used in Topping et al. 2015. See text for further detail.

For the present project, the land cover map was assembled using the following rules: (1) The field layer was the bottom layer and (2) the lake and streams (freshwater theme) was put on top of the field layer. (3) Dry natural areas (nature theme) were then added, but only to cells not already occupied by 1 or 2. (4) Similarly, built-up areas (builtup theme) were added to cells not already occupied by 1, 2 or 3. (5) Wet natural areas (wetnature theme), (6) Cultural features (cultural theme), (7) roads (road theme) & (8) sea (raster layer landsea) were then added sequentially onto preceding map. Finally (9) buildings (raster layer buildings_250) were added. After this process there may still number of cells without land cover type, depending of the quality of the input data. There may be several strategies to complete the map: One option is to compare gaps to a recent orthophoto and manually assign a land-use category, but this is a very time-consuming for larger maps. Another option is to fill gaps with a randomly chosen landscape feature (stratified to represent the general landscape structure). For the present study we used an existing, older land cover map of Denmark (AIS, Stjernholm et al. 2010) to fill in gaps.

#### Preparing maps for ALMaSS simulation

##### Convert land cover map to ALMaSS landscape

Finalizing the ALMaSS landscape is done in the fifth and last section of the python script (Finalize). The land cover map contains more detail than are used in ALMaSS for most applications. This information has been retained up until now, but will need to be condensed into the landscape element types to be used in ALMaSS. This is a simple reclassification based on a text file with the landscape element codes used in ALMaSS. All features in the raw ALMaSS landscape, both features consisting of single or of multiple raster cells, have a unique value that is common to all cells within the feature. This is achieved by regionalizing (Fig. 8) the raster before exporting the map as an ASCII file. This ASCII file is then converted to a binary format whereby all cells are represented by a 32-bit integer, in ALMaSS designated as a landscape binary file (.lsb). This step may not be necessary for other applications, but if so the LSBConverter.exe program can be obtained from the ALMaSS project site on CCPForge http://ccpforge.cse.rl.ac.uk/gf/project/almass

**Figure 8.**
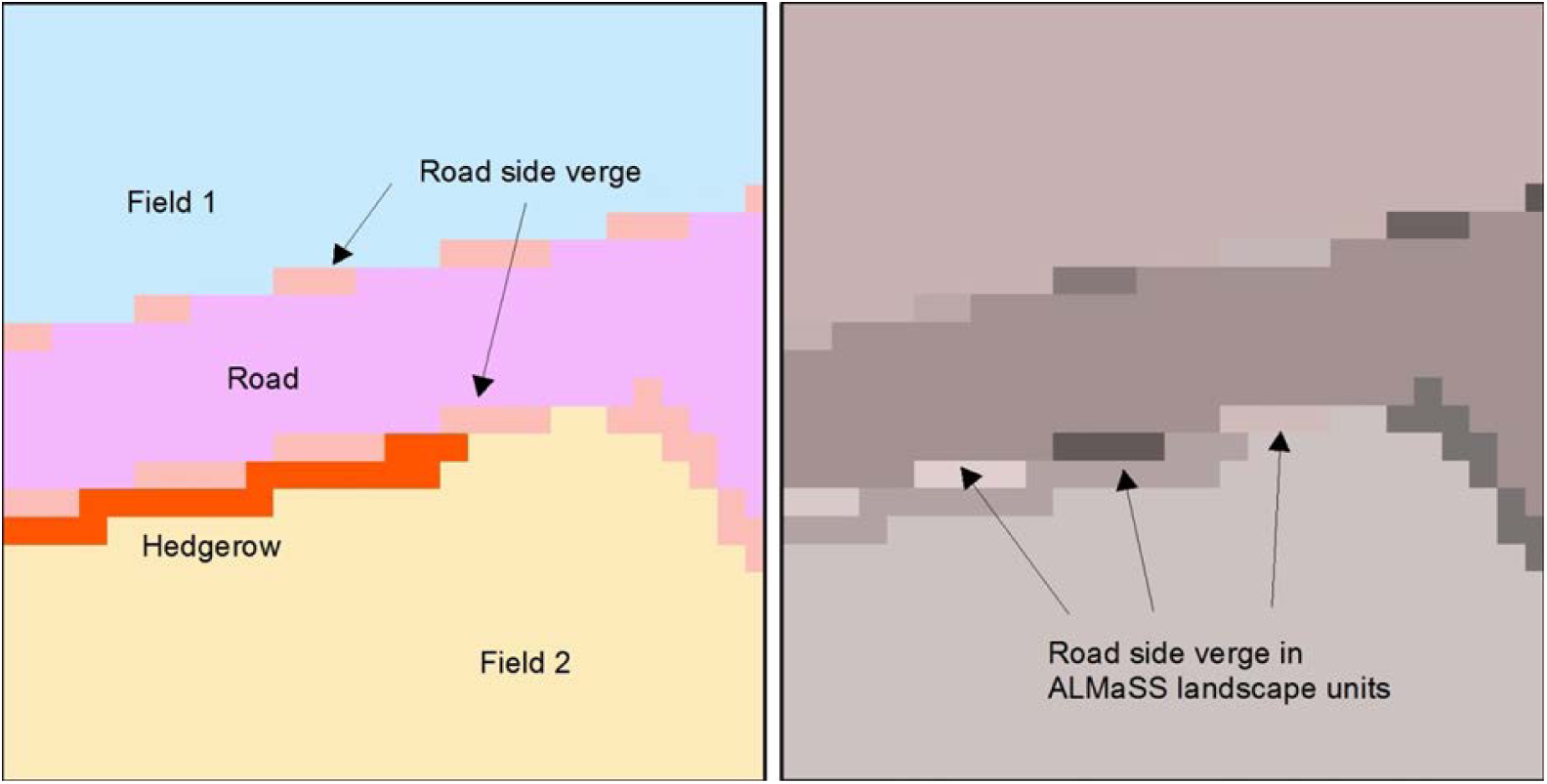
Regionalization. This figure shows how a reclassified land cover maps is transformed into an ALMaSS model landscape. ALMaSS uses landscape units (defined as clusters of pixels of the same land cover type) to store landscape information. Each unit has a unique identifier (as a polygon would) and a land cover code. This is achieve by the regionalize function in ArcGIS. The map on the left shows the classified land cover map (with two fields, a road, road side verge and a hedgerow. The map on the right shows the results of the regionalization. The two fields, the road and the hedgerow are uninterrupted and treated as units. The road side verge, on the other hand, is fragmented and each fragment must be treated as a unique ALMaSS landscape unit.

##### Polygon reference file

Each polygon on the final ALMaSS landscape only contains one value which is the unique ID of the polygon. All additional information about the polygon, such as landscape element type and farm ownership is contained in the polygon reference file. The polygon reference file is a text file containing a unique ID on all polygons in the landscape, the landscape element type of each of the polygons, the number of cells belonging to each polygon, a reference indicating farm ownership and optionally the soil type of each polygon (see Table 2). Upon loading the polygon reference file in ALMaSS, the program adds coordinates for the polygon centroid in the coordinate system used in the simulation. Additionally, columns with openness scores (used when modelling geese) and a unique ID for unsprayed field margins on fields are added.

**Table 2.** Table showing the structure of the polygon reference file when first imported to ALMaSS. The first line is indicating the number of lines in the file (here just an example, the actual file will contain many more lines), second line the headers and all following lines the actual values. The value -1 is used to indicate NA.

| PolyType | PolyRefNum | Area | FarmRef | UnSprayedMarginRef | SoilType |
| --- | --- | --- | --- | --- | --- |
| 80 | 0 | 2004 | -1 | -1 | -1 |
| 110 | 1 | 8105 | -1 | -1 | -1 |
| 94 | 2 | 163599 | -1 | -1 | -1 |
| 110 | 3 | 3 | -1 | -1 | -1 |
| 123 | 4 | 278954 | -1 | -1 | -1 |
| 110 | 5 | 62 | -1 | -1 | -1 |
| 20 | 6 | 43825 | -1 | -1 | 5 |

##### Farm reference file

ALMaSS needs a farm reference file for the each simulated landscape. The file is a text file with two columns, one being the farm reference number (a unique ID for each farm in the landscape) and a column with the farm type of each of the farms (see “Farm classification”).

##### Making the reference files

To create the polygon reference file, the attribute table from the final land cover map needs to be exported manually from ArcGIS. With this the attribute table and information about farm ownership (which farm owns which fields) a minimal polygon reference file can be made. If soil type information is available it can be used to improve modelling of growth of vegetation in the simulation. The task is to merge these three pieces of information together. Merging can be done in any standard data base program or programming language and the optimal choice of tool will depend on the format in which farm- and soil type information is available. To prepare the polygon- and farm-reference files for Topping et al. 2015 we used R and functions in the R packages ralmass (Dalby 2015), devtools (Wickham & Chang, 2015) and data.table (Dowle et al. 2014). An example script is provided below.

**Figure 9.**
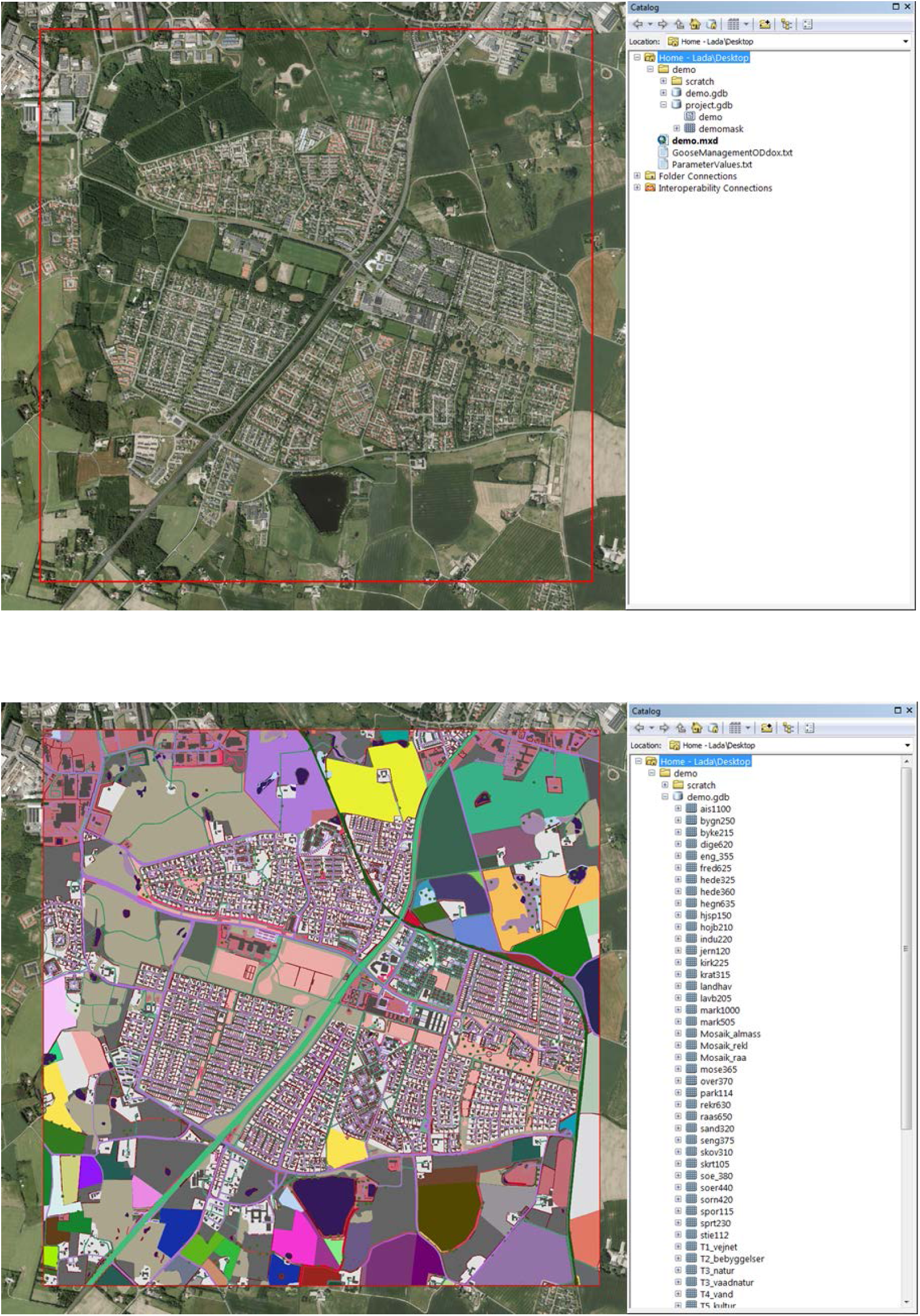
Study area outline and final map. In (a - top) the red outline show the area to make a simulation landscape for. The ortophoto is showing a recent image of the actual landscape. In the catalog pane on the right hand side of (a) is showing the file structure for the workspace that is needed to run the python script. A Scratch folder to store temporary files, a geodatabase (project.gdb) holding the vector outline (demo, shown in red on in the map view) and the raster version of this outline (here name demomask) and finally a geodatabase to store outputs from the script (here named demo.gdb). In (b - bottom) the final map is shown. In the catalog pane on the right hand side the demo geodatabase is shown with all its containing layers expanded (some not shown). These are the individual raster layers, the thematic maps and the final maps. They are stored to enable quality control of the individual steps in the process after the final map is made.

R script

Set-up and import the attribute table from ArcGIS.

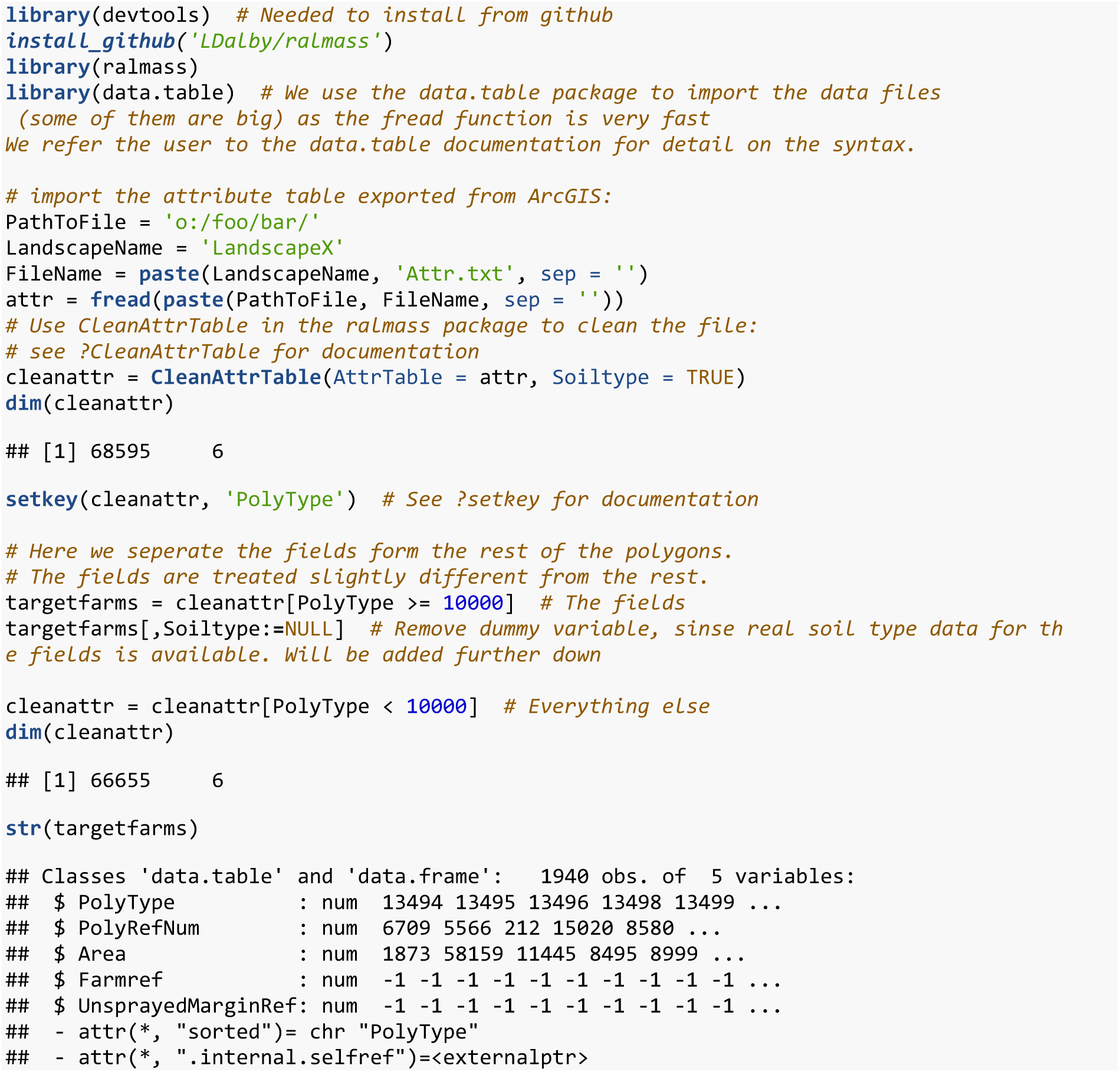

Next we read in the farm information. In this example the data is stored in a text file where each field is a row in the data set. Each field has a unique ID for the farm to which it belongs.

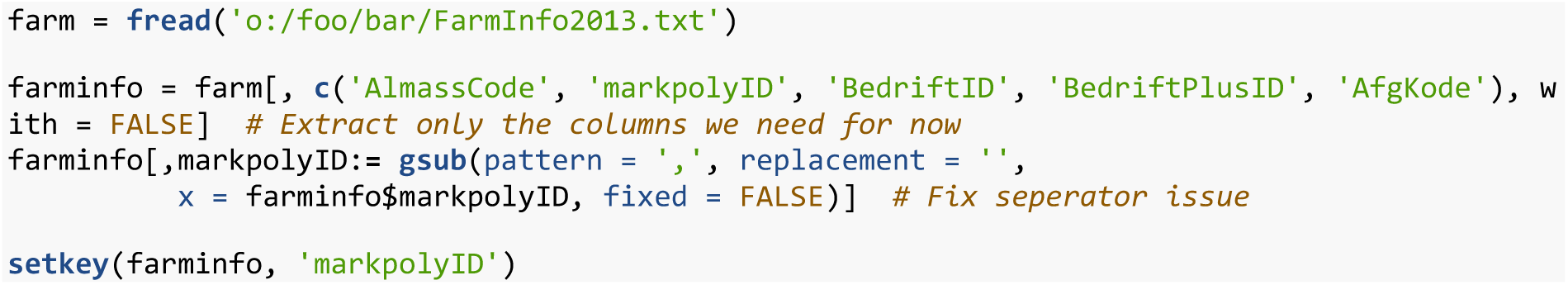

In this particular case we do have soil type information for the fields, so we load that.

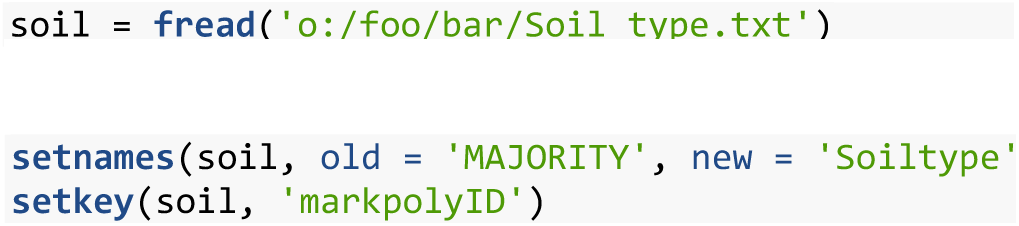

With all three pieces of information we merge the datasets using the unique field polygon ID as key.

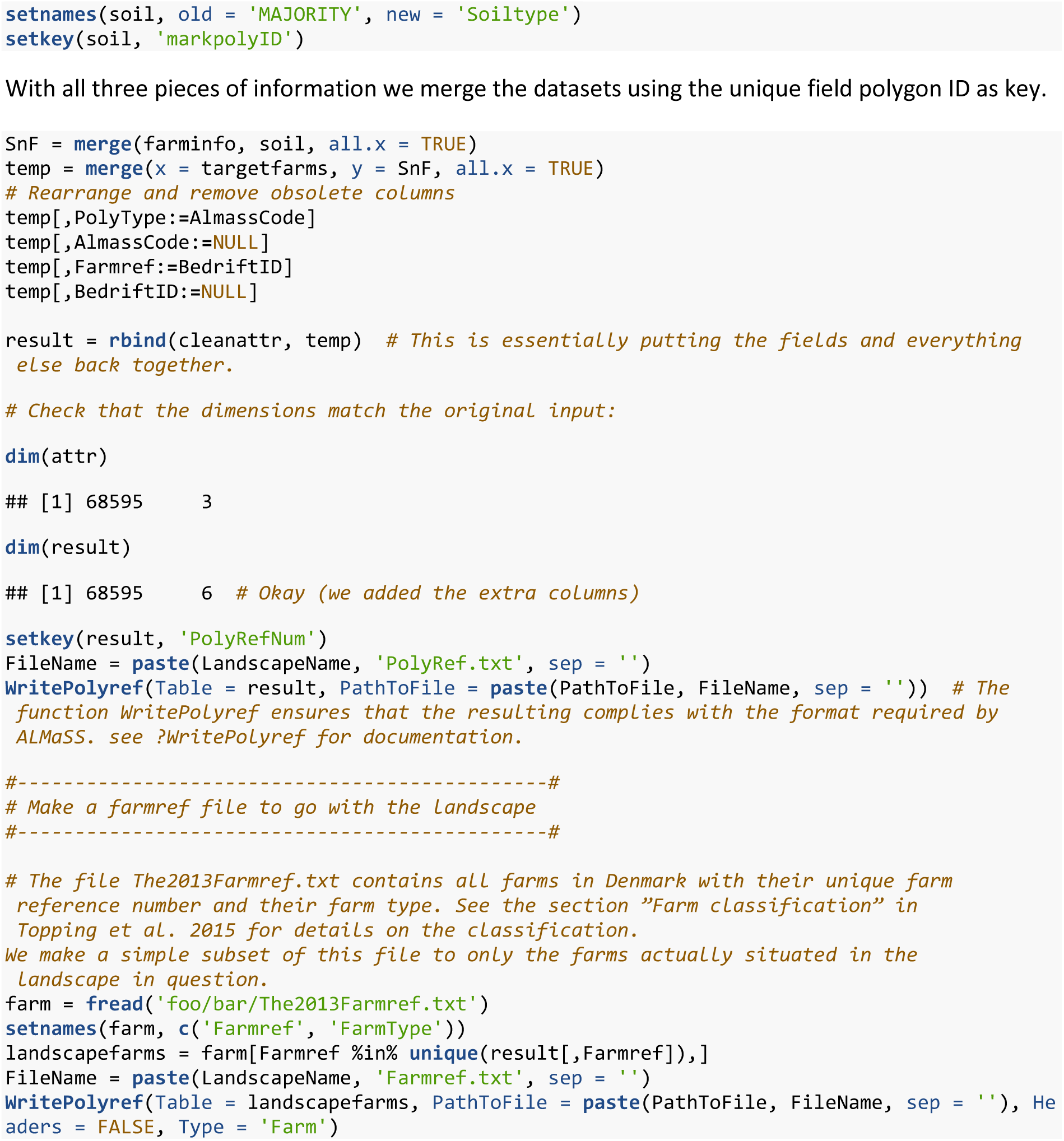

## Appendix 2: overview of ALMaSS farm management

Aspects of the farm management have been described in a number of ALMaSS publications separately (incl. Topping, Ostergaard et al. 2003, Topping, Hansen et al. 2003, Thorbek and Topping 2005, Topping and Olesen 2005, Topping 2011, Parry, Topping et al. 2013), and are described in the program documentation ODdox (Topping, Hoye et al. 2010) format (e.g. Topping 2009),but have not been described as a whole in text form. Since much of the power of ALMaSS for use in environmental risk assessment comes from its ability to handle detailed farm management, a general overview of the processes is provided here.

### Purpose

The farm management in ALMaSS creates a dynamic and emergent pattern of both crop coverage patterns at landscape and field scales but also patterns of farm management activities in time and space. This information is available via the ALMaSS Landscape class interface for any object in the ALMaSS simulations. Thus the farming module’s purpose is to simulate farming realistically at landscape and farm scales and to provide information on vegetation changes and farming activities to the Landscape class.

### Class structure (class names in italics)

The overall class handling environmental information in ALMaSS is the *Landscape.* This class contains a map of the landscape represented by homogenous polygons *(LE* (landscape element)) classified into types. Those designated as type field are represented by the *Field* class, and each is linked to the farm that manages it (either based on real information e.g. GLR (see Methods), or specified as the user wishes). All farms are represented by the *Farm* class, and all farms are held in lists managed by the class *FarmManager,* which is instantiated as a class member of *Landscape.* Crops grown on a field are also classes, each is a specific descendent class of the main *Crop* class, e.g. *SpringBarley.* These classes include the implementation of the specific crop husbandry for that crop.

### Process overview

This version of the methods assumes the use of the *Farm* class rather than the *OptimisingFarm* class. The *Farm* class cannot really be considered an agent, since they have no goals on which to base decisions but act following predefined rules made variable by the introduction of stochasticity as probabilistic rules. In contrast *OptimisingFarm* objects are true agents and have goals and more flexible decision and learning behaviour. *Farm* class is the most common usage of ALMaSS since it requires much less information to set up compared to the more complex alternative.

At the highest level of organisation data is used to classify farms (see manuscript section 2.1.2), which in turn determines the rotation used by all farms. This information is used, together with the mapped field polygons and their associated farm classified into types, as the basis for determining crop coverage by area at the landscape and farm level. Day-to-day farm management determines the actual cover on fields and in the landscape, crop structure in terms of height and green and total leaf-area index, and farming activities (e.g. ploughing) (Fig. 1).

**Figure 1:**
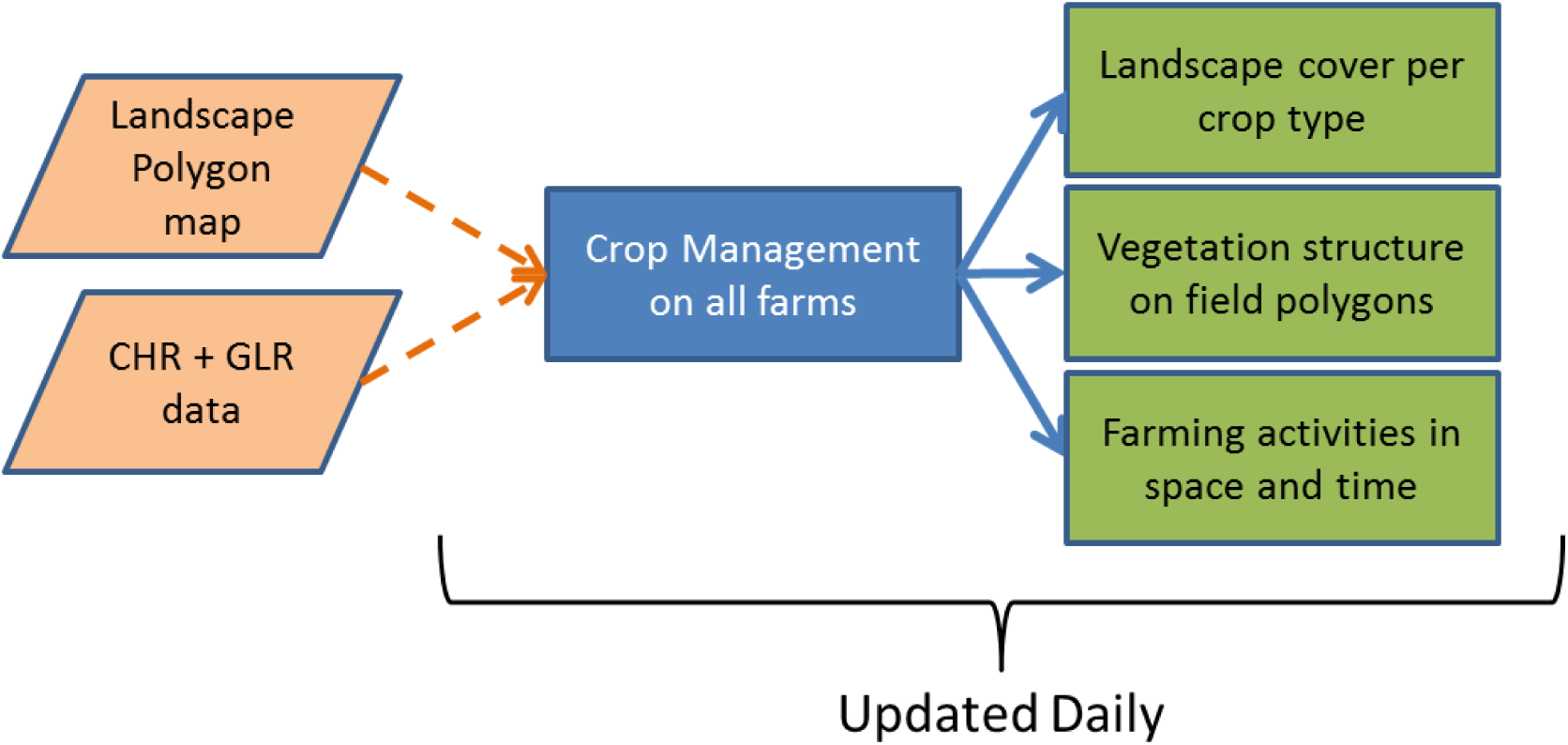
The farm management inputs and outputs at the general level of organisation

On a daily basis the management is carried out at field polygon level and comprises a crop husbandry model fed by system data inputs, linked to a crop growth model and jointly creating outputs at the field polygon level (Fig. 2).

**Figure 2:**
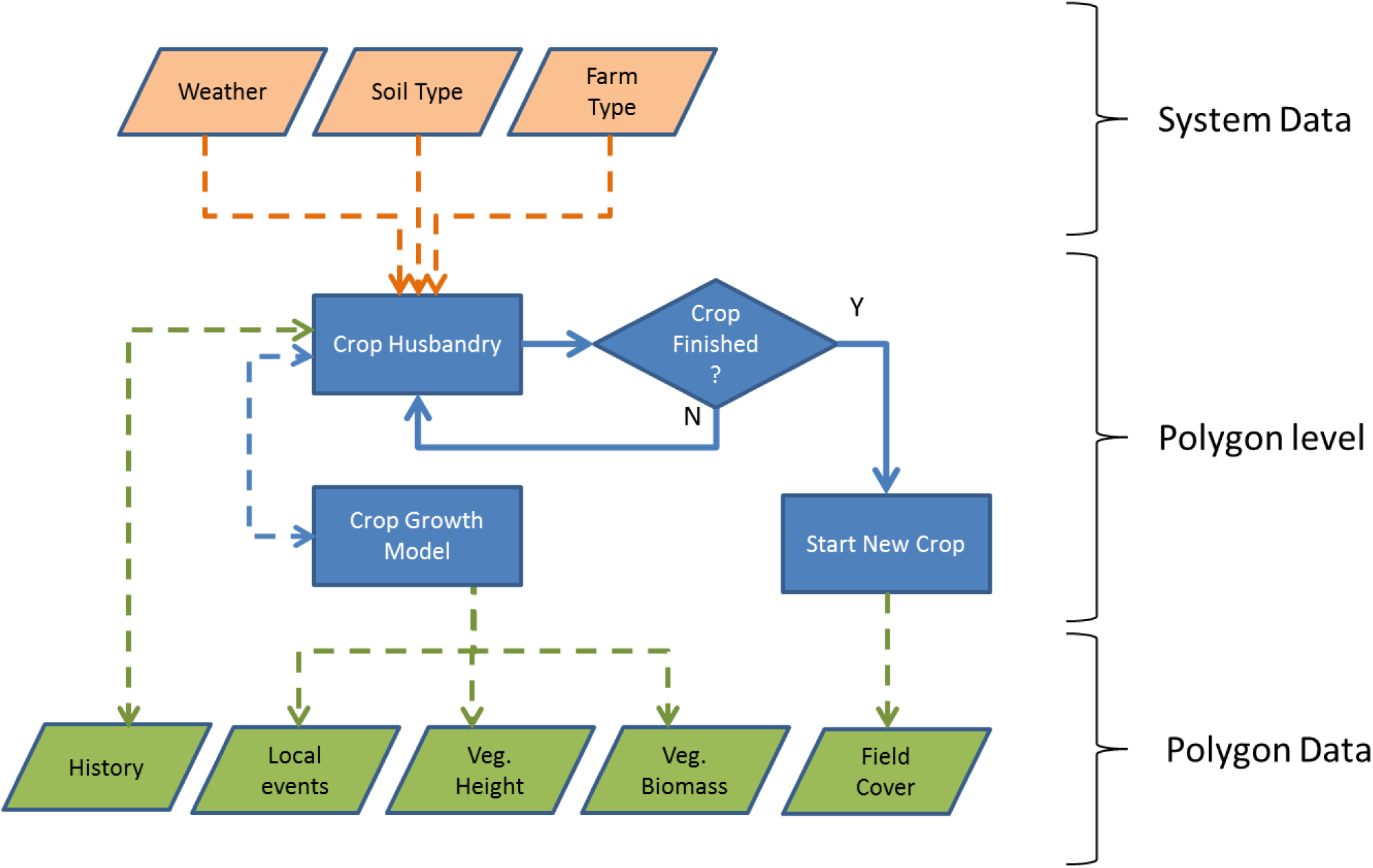
Data flows into and out of farm management at the field polygon level. Dashed arrows are data flows, solid arrows process flow

The basis for the crop growth model is described by (Topping and Olesen 2005). This results in a series of growth phases with height, leaf-area total, and leaf area green as a function of summed day degrees from the start of the growth phase (e.g. Table 1). Using these growth phases it is possible to recombine them in different orders to represent all Danish crops e.g. an autumn sown crop will get ‘From Sowing’, ‘From Jan 1^st^’ ‘From Mar 1^st^’, ‘From harvest 1’, whereas a spring crop might be ‘From mar 1^st^’, ‘From Sowing’. ‘From Harvest 1’. Management e.g. harvest or sowing can cause a change in growth phase and a sudden change in vegetation characteristics (e.g. height after harvest), hence at the start of each growth phases it is possible (but not obligatory) to set the vegetation characteristics to a particular value. For each growth phase the rates of change for the three response variables per day degree (calculated between inflection points on the curve, i.e. rows in Table 1) are stored as vegetation specific growth curves. These specific change rates are applied on a daily basis (Fig. 3) and vegetation characteristics updated based on the leaf-area index and height changes (cover and biomass can be calculated from these e.g. using Beer’s Law for cover).

**Table 1:** An example of the format of the vegetation growth curve data used in ALMaSS as input to the vegetation daily growth model showing the five growth phases possible.

| Growth Phase | TSum | LAI-Total | LAI-Green | Height |
| --- | --- | --- | --- | --- |
| **From Jan 1st** | 0 | 4.83 | 0 | 70 |
|  | 105 | 3.85 | 0 | 56 |
| **From Sowing** | 99999 | 0 | 0 | 0 |
| **From Mar 1st** | 0 | 3.85 | 0 | 56 |
|  | 289 | 4.5 | 1 | 65 |
|  | 695 | 5 | 1.25 | 73 |
|  | 1017 | 5.5 | 0 | 80 |
|  | 2619 | 5.5 | 0 | 80 |
|  | 2787 | 4.83 | 0 | 70 |
|  | 99999 | 4.83 | 0 | 70 |
| **After cutting/harvest 1** | 0 | 0.3 | 0 | 10 |
|  | 289 | 4.5 | 1 | 65 |
|  | 695 | 5 | 1.25 | 73 |
|  | 1017 | 5.5 | 0 | 80 |
|  | 2619 | 5.5 | 0 | 80 |
|  | 2787 | 4.83 | 0 | 70 |
|  | 99999 | 4.83 | 0 | 70 |
| **From cutting/harvest 2** | 99999 | 0 | 0 | 0 |
|  | 289 | 4.5 | 1 | 65 |
|  | 695 | 5 | 1.25 | 73 |
|  | 1017 | 5.5 | 0 | 80 |
|  | 2619 | 5.5 | 0 | 80 |
|  | 2787 | 4.83 | 0 | 70 |
|  | 99999 | 4.83 | 0 | 70 |

**Figure 3:**
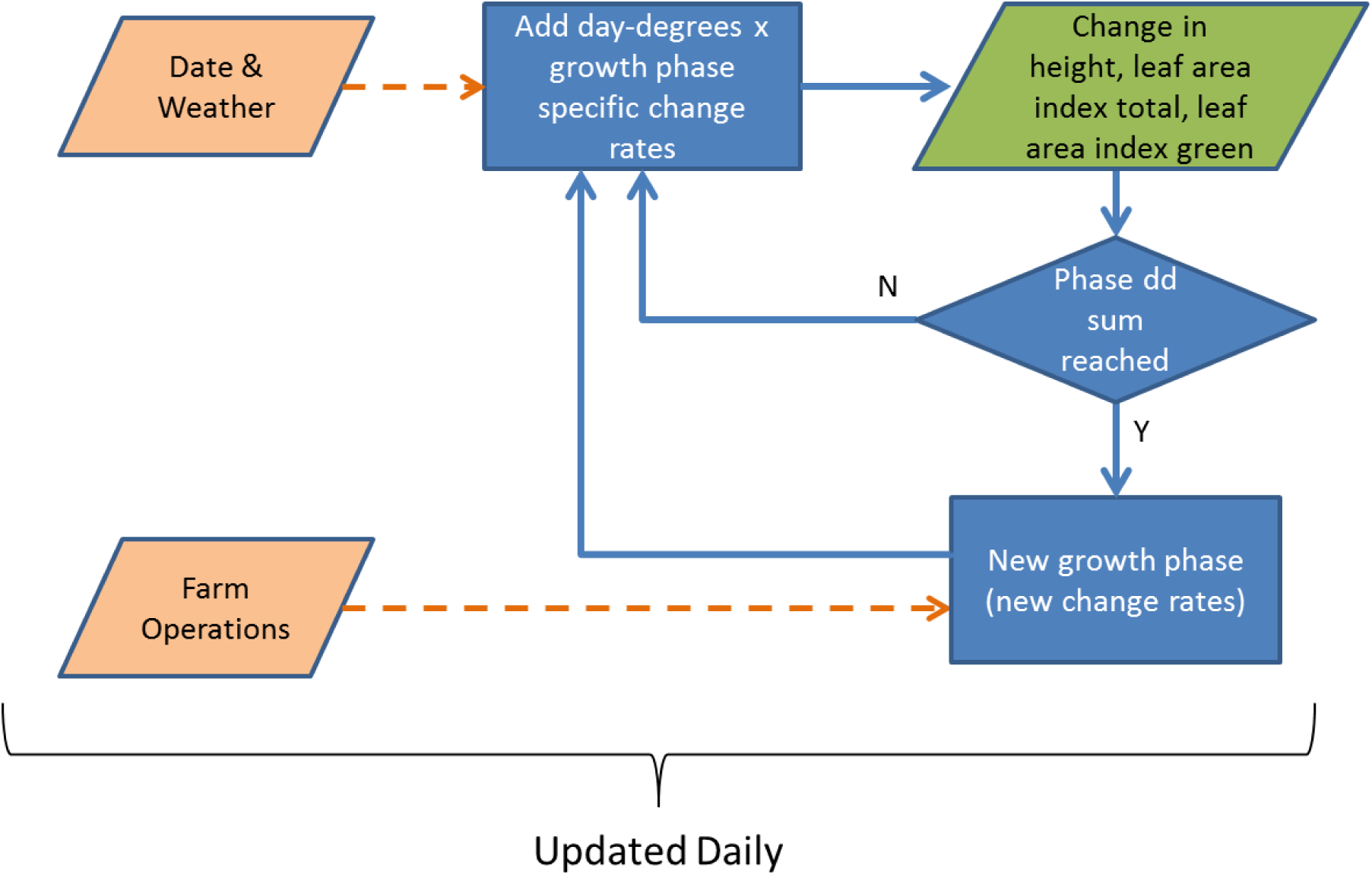
The crop growth model data and process flows leading to changes in vegetation height and leaf area index. Data is represented by dashed lines, process flows by solid lines

Crop husbandry results in recording the farm operation events that occur on each field each day. The sequence of events and their conditions is specified by a unique crop management plan for each crop. Examples of the implementation can be found in the ALMaSS ODdox (e.g. Topping 2009). These plans are long and complicated and hence only a section of a representative plan is shown here to demonstrate the process (Fig. 4). Whether an operation is carried out on a particular day is determined by a probability distribution resulting in a spreading of operations in time within the permitted period of action (start to end date inclusive in Fig. 4). It is also dependent upon weather or history events e.g. spraying a second herbicide may only happen if the first was sprayed and only under low wind-speed and no precipitation conditions. In this way real agronomic constraints can be included.

Some constraints for farm operations are programmed into the farm class and are thus general to all attempts to carry out the operation e.g. weather constraints to pesticide sprays; others are specific to the crop husbandry plan and the actual polygon it is being applied to, e.g. history of operations.

**Figure 4:**
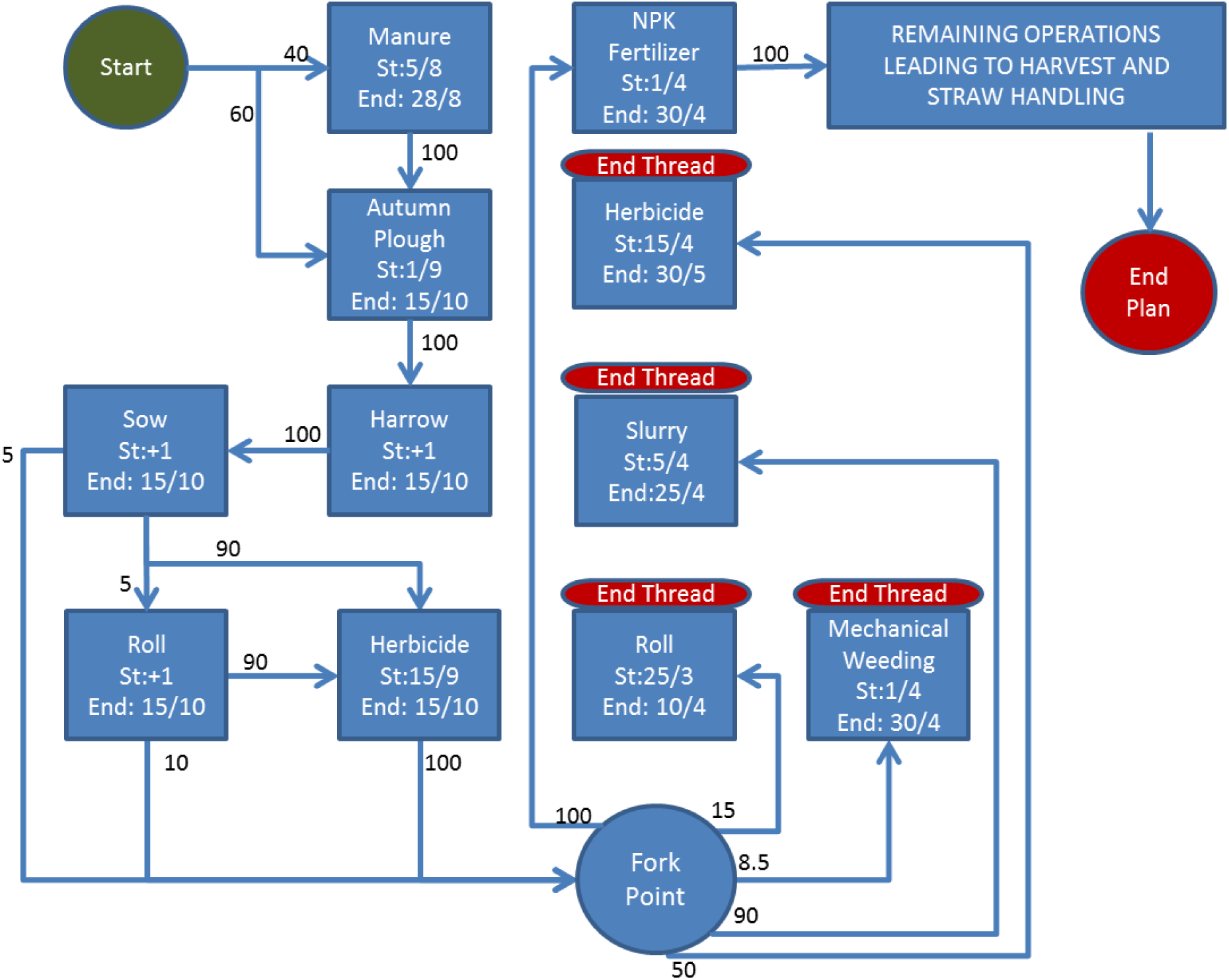
Flow diagram showing events in the husbandry plan for triticale up to end of April. Black numbers are base % making the choice to carry out an operation (blue boxes). Note threads fork from the ‘Fork Point’ and carry on parallel actions until each thread ends. One thread, in this case the NPK thread carries on to the rest of the husbandry plan (not shown). St = start date End = end date for an operation

## Appendix 3: 10 Danish ALMaSS Landscapes

We here present a visual overview of each of the 10 Danish landscapes used in Topping et al. 2015 along with a map of Denmark with each of the landscapes shown. Note that at the scale used here it is not possible to map all landscape elements present in the landscapes. See Appendix 1 in Topping et al. 2015 for full details.

Topping, C. J., Dalby, L. & Skov, F 2015: Landscape structure and management alter the outcome of a pesticide ERA: evaluating an endocrine disruptor using the ALMaSS European Brown Hare model

**Figure 1:**
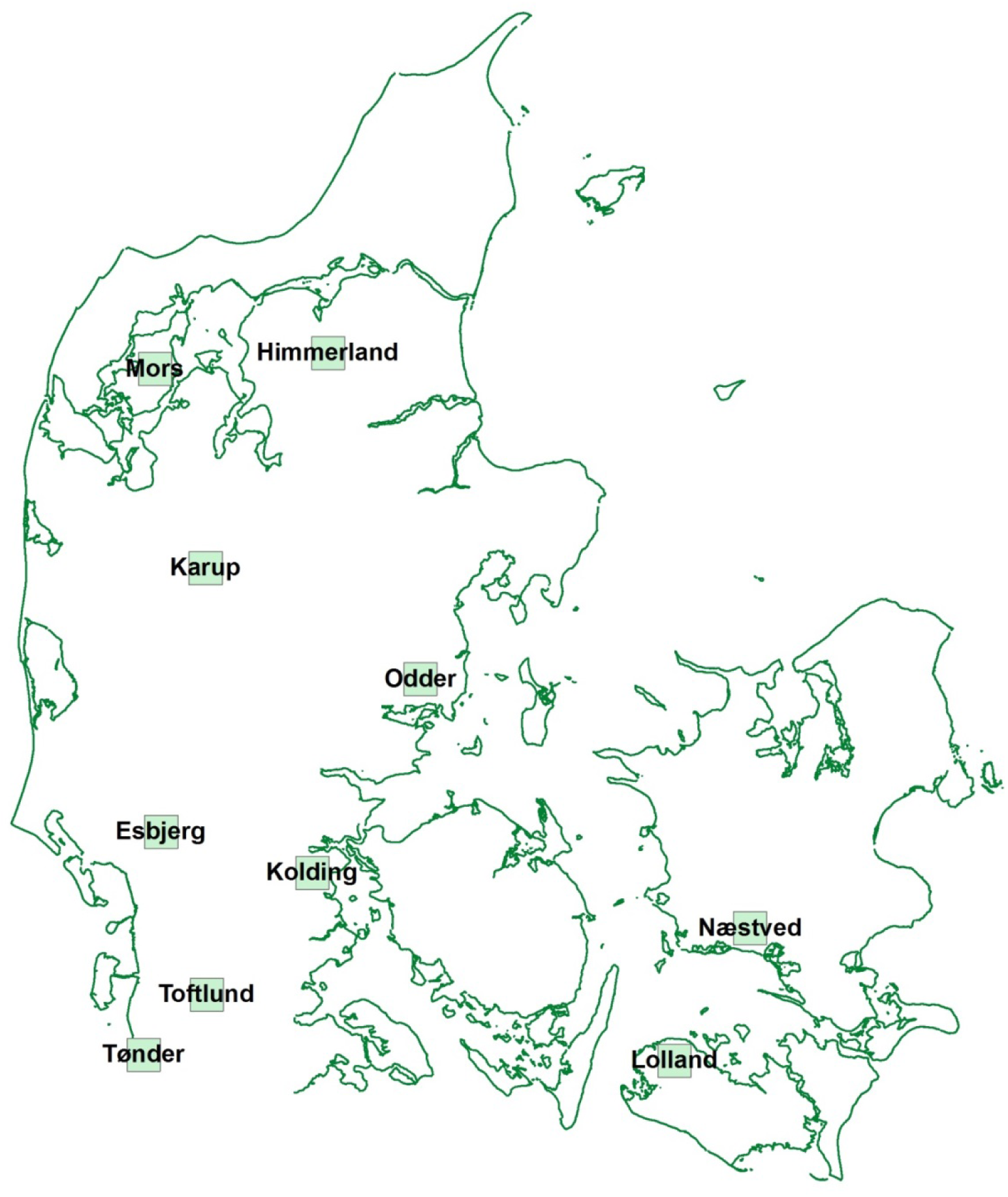
The location of each of the 10 landscapes in Denmark

**Figure 2:**
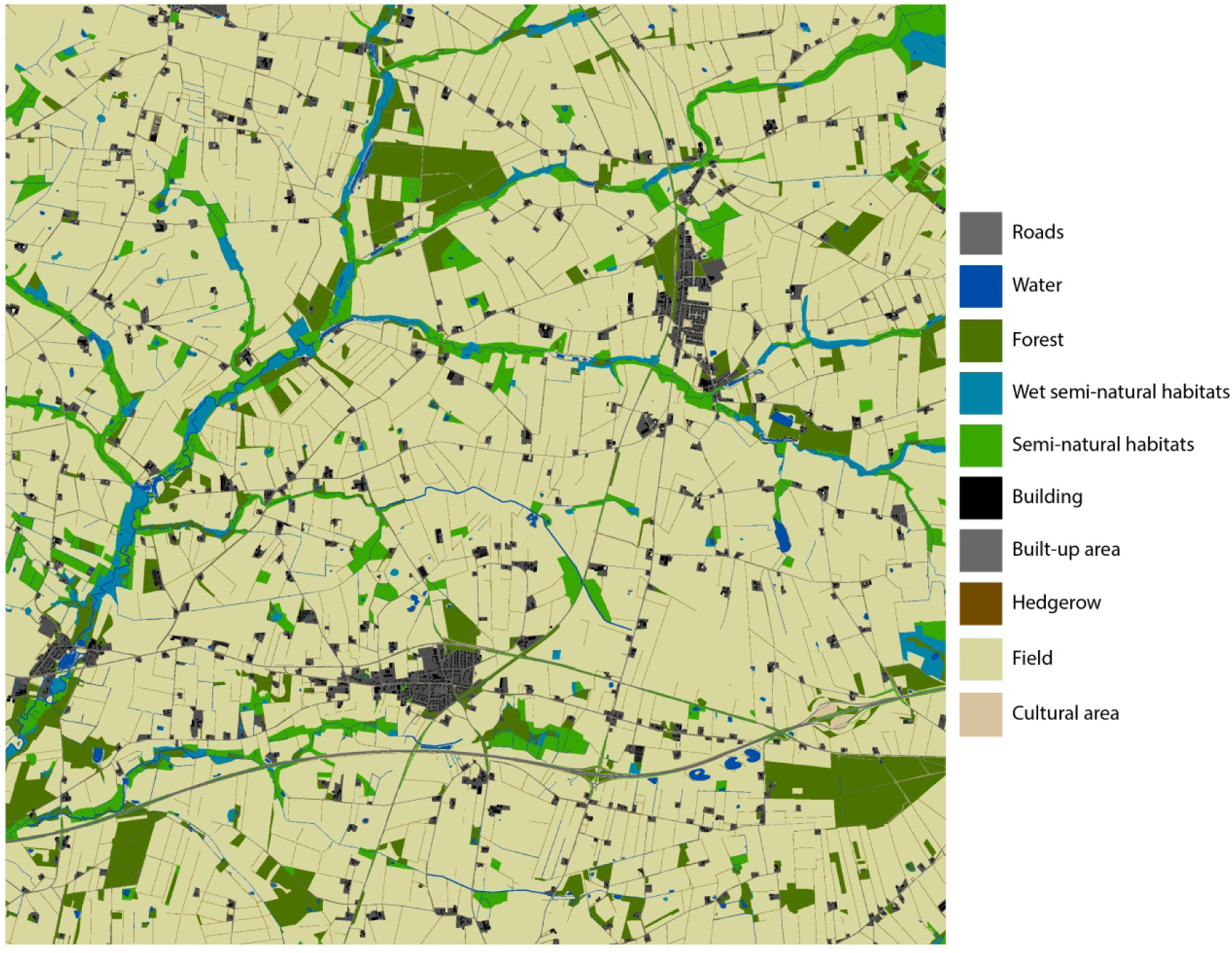
Esbjerg

**Figure 3:**
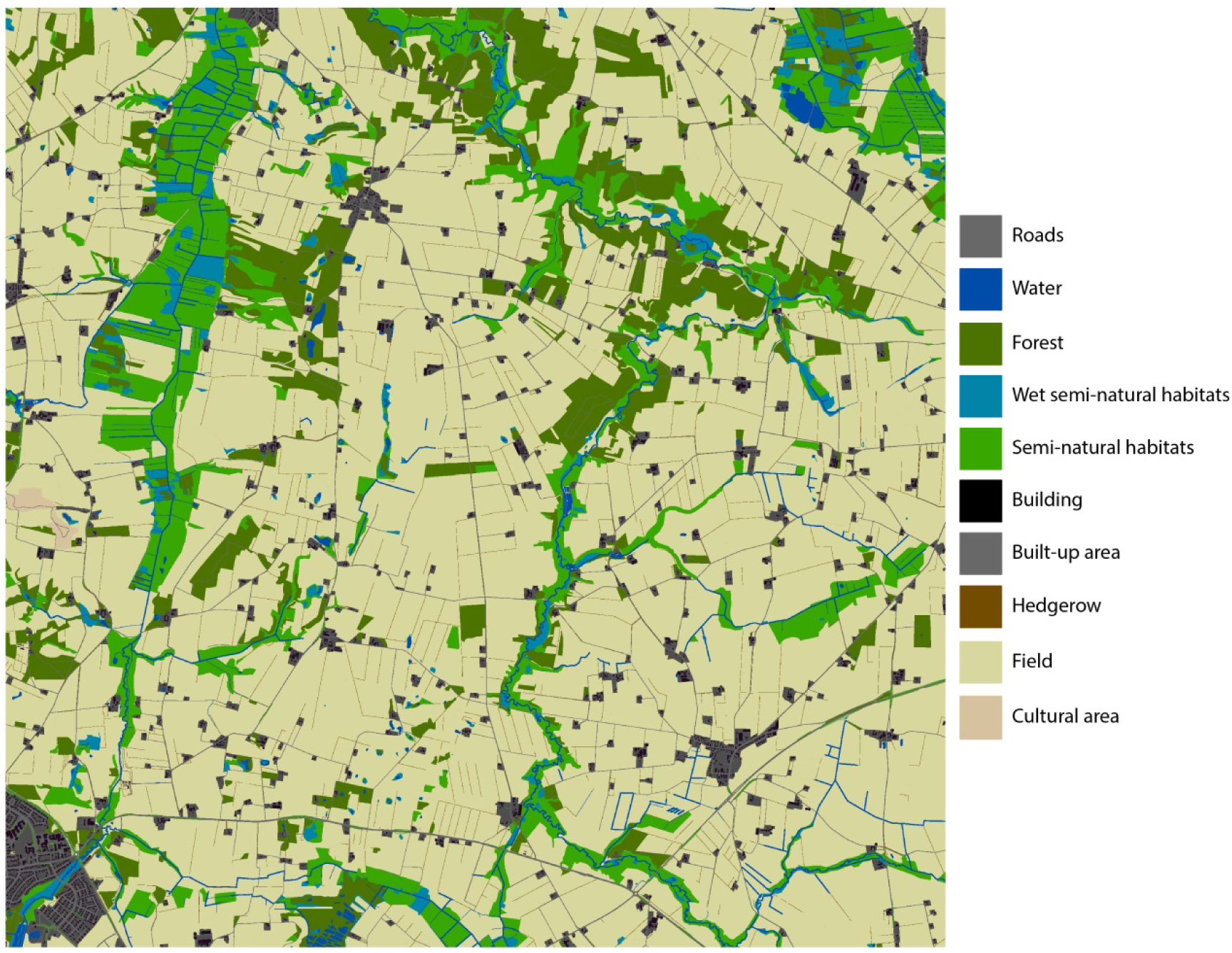
Himmerland

**Figure 4.**
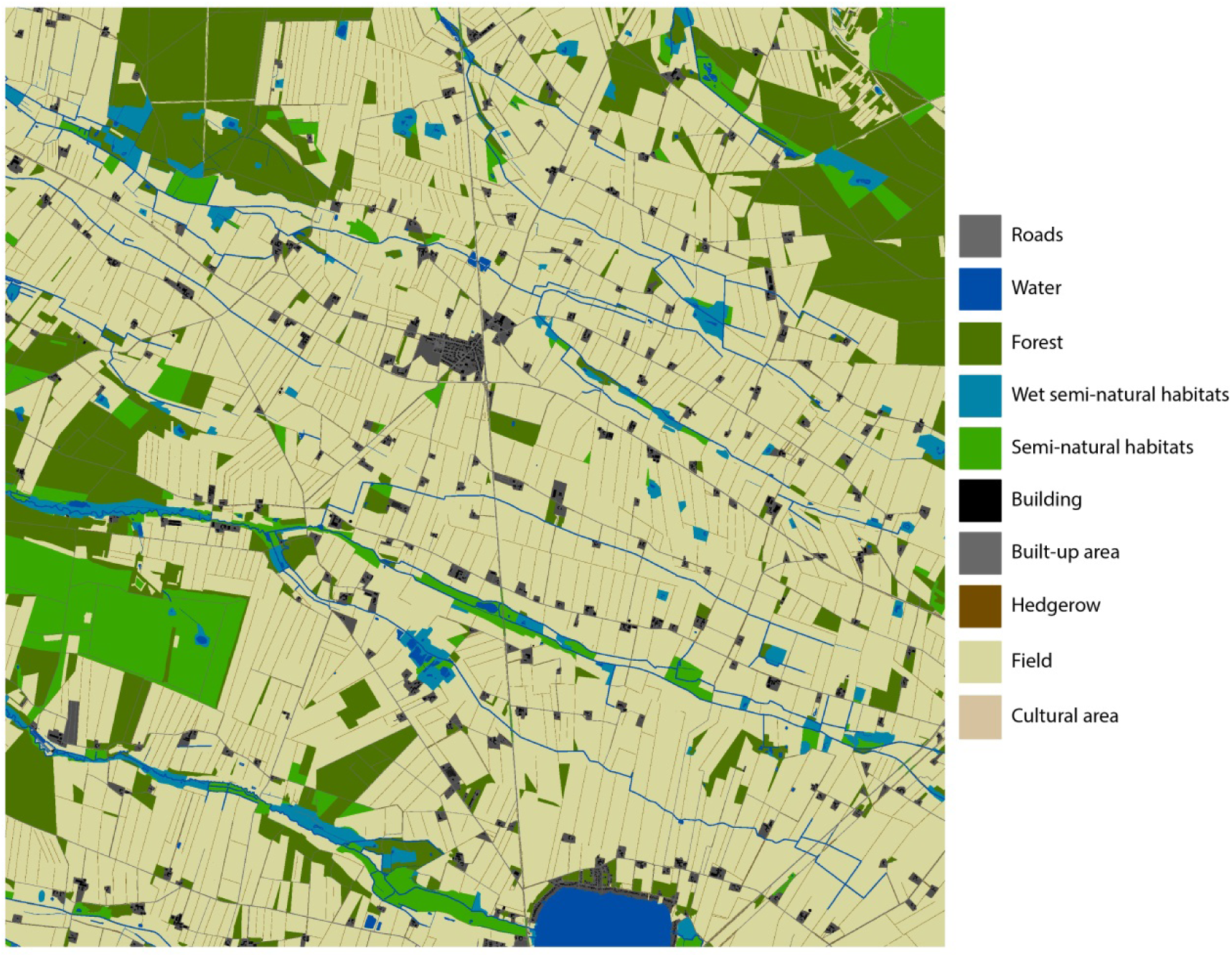
Karup

**Figure 5.**
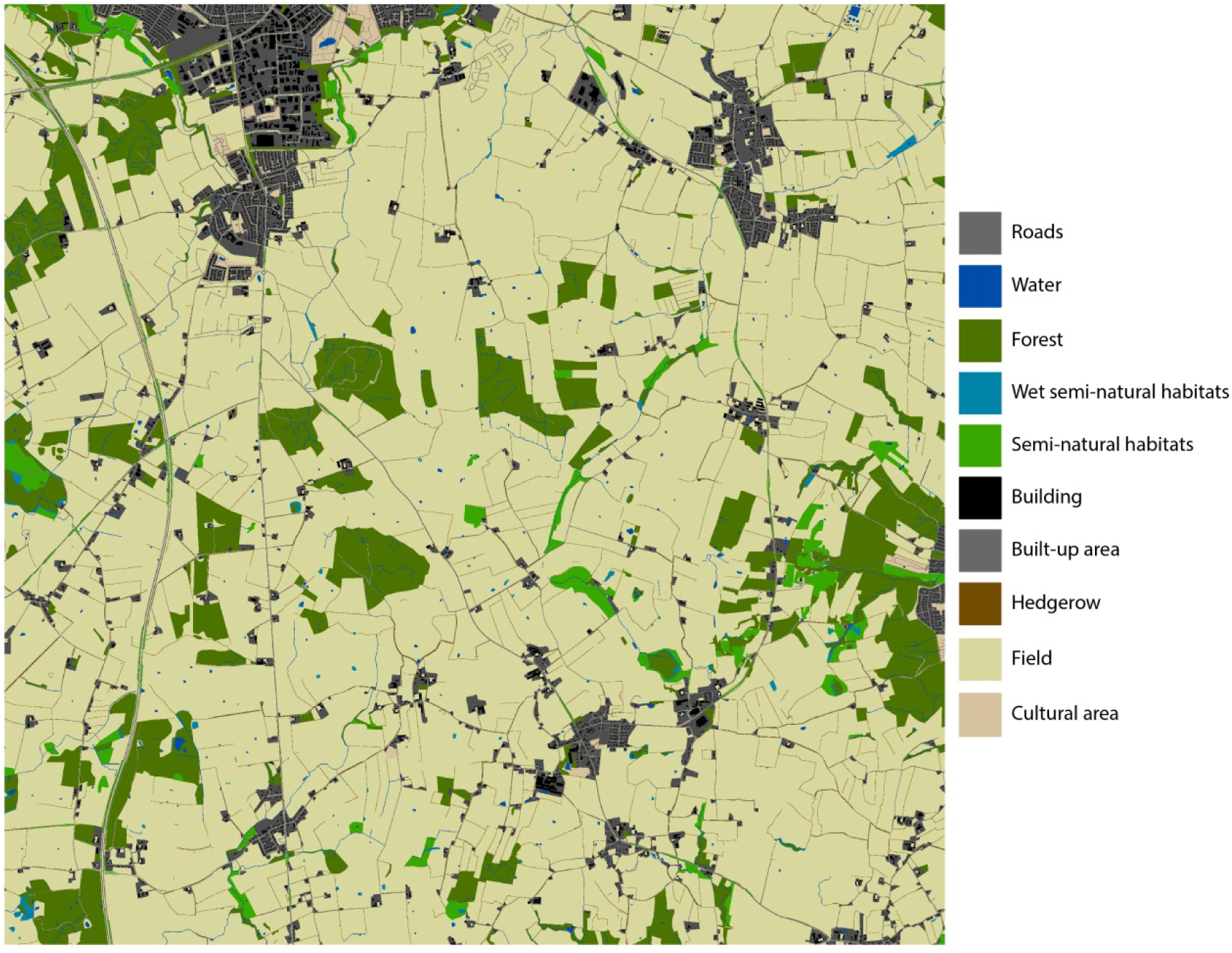
Kolding

**Figure 6.**
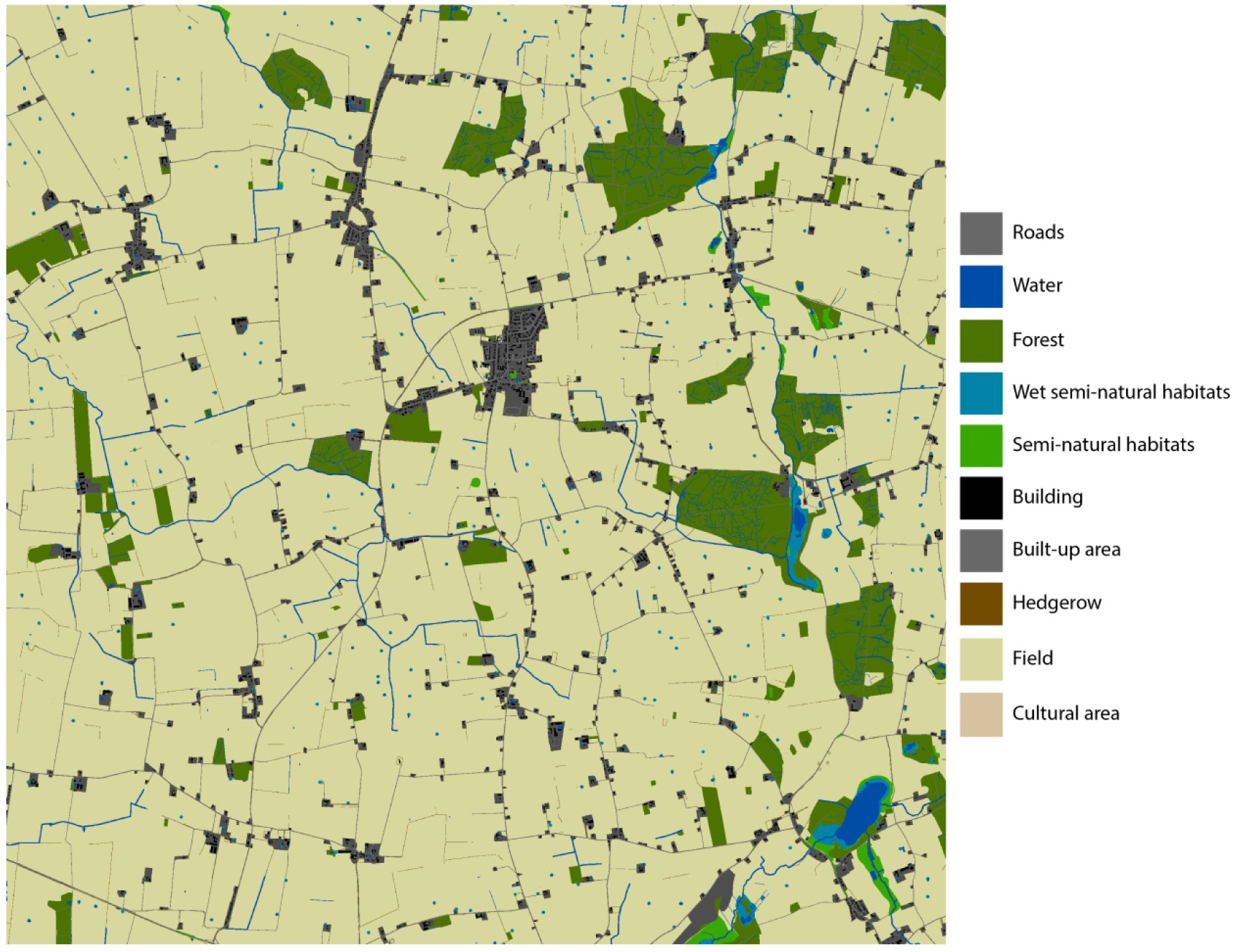
Lolland

**Figure 7.**
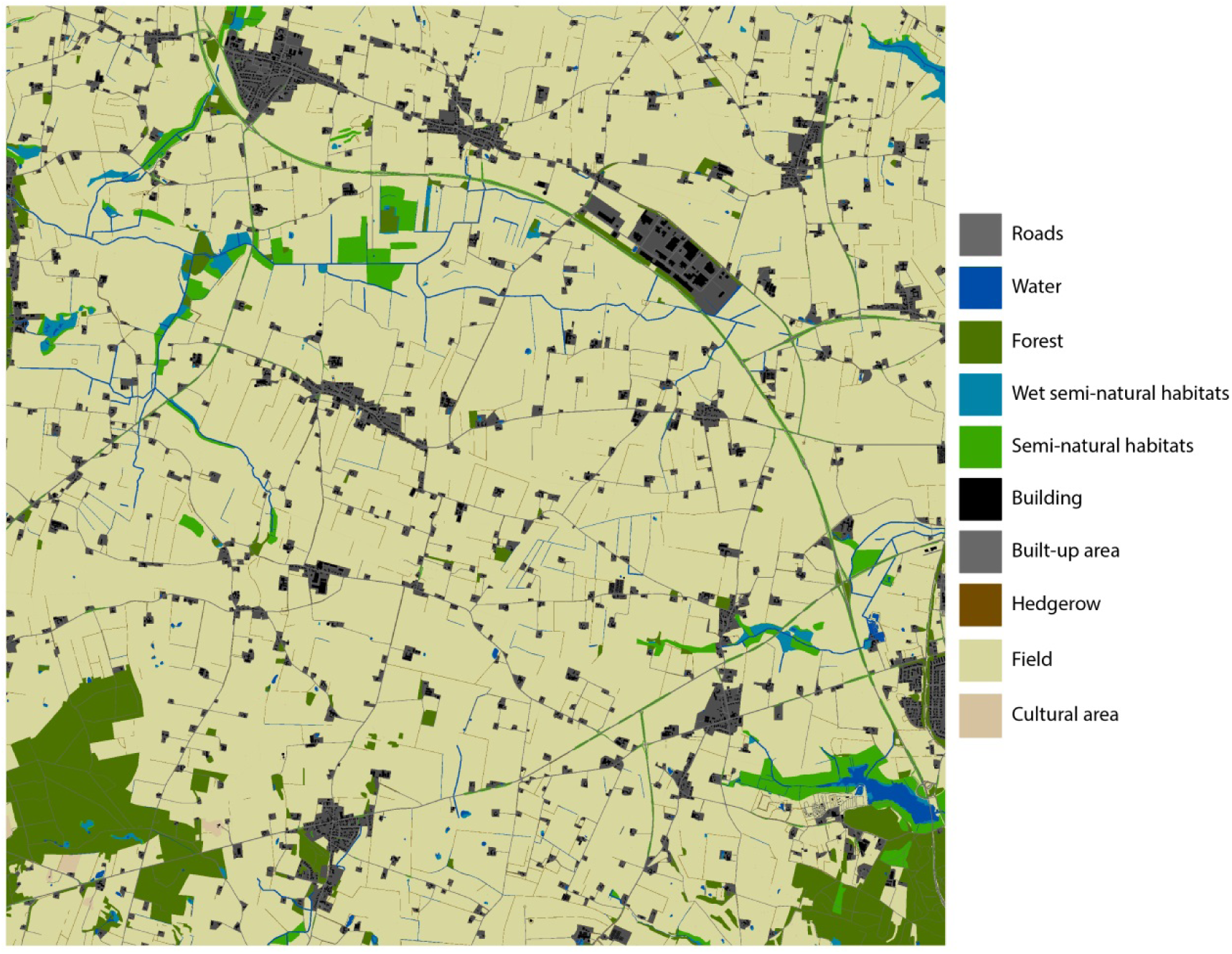
Mors

**Figure 8.**
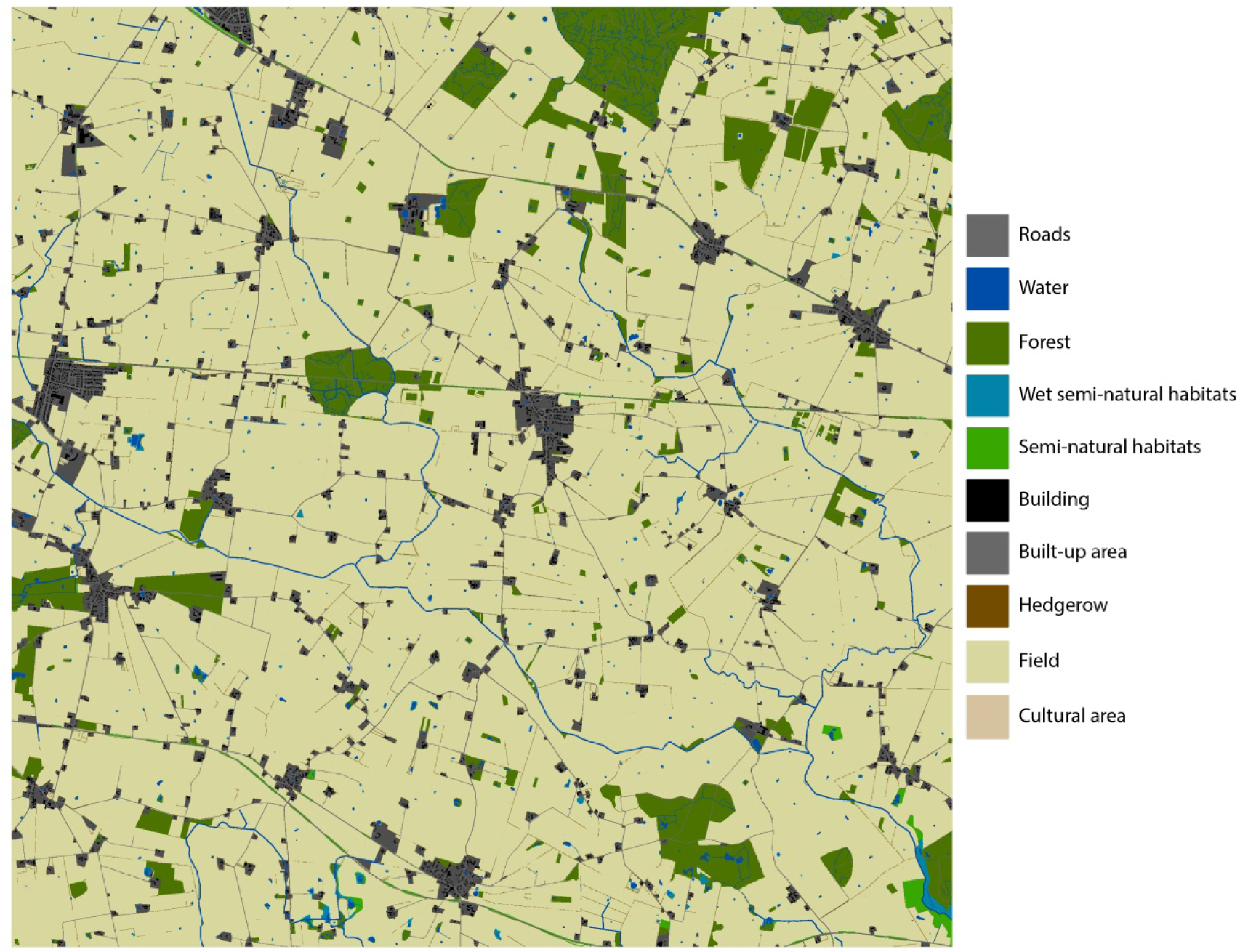
Næstved

**Figure 9.**
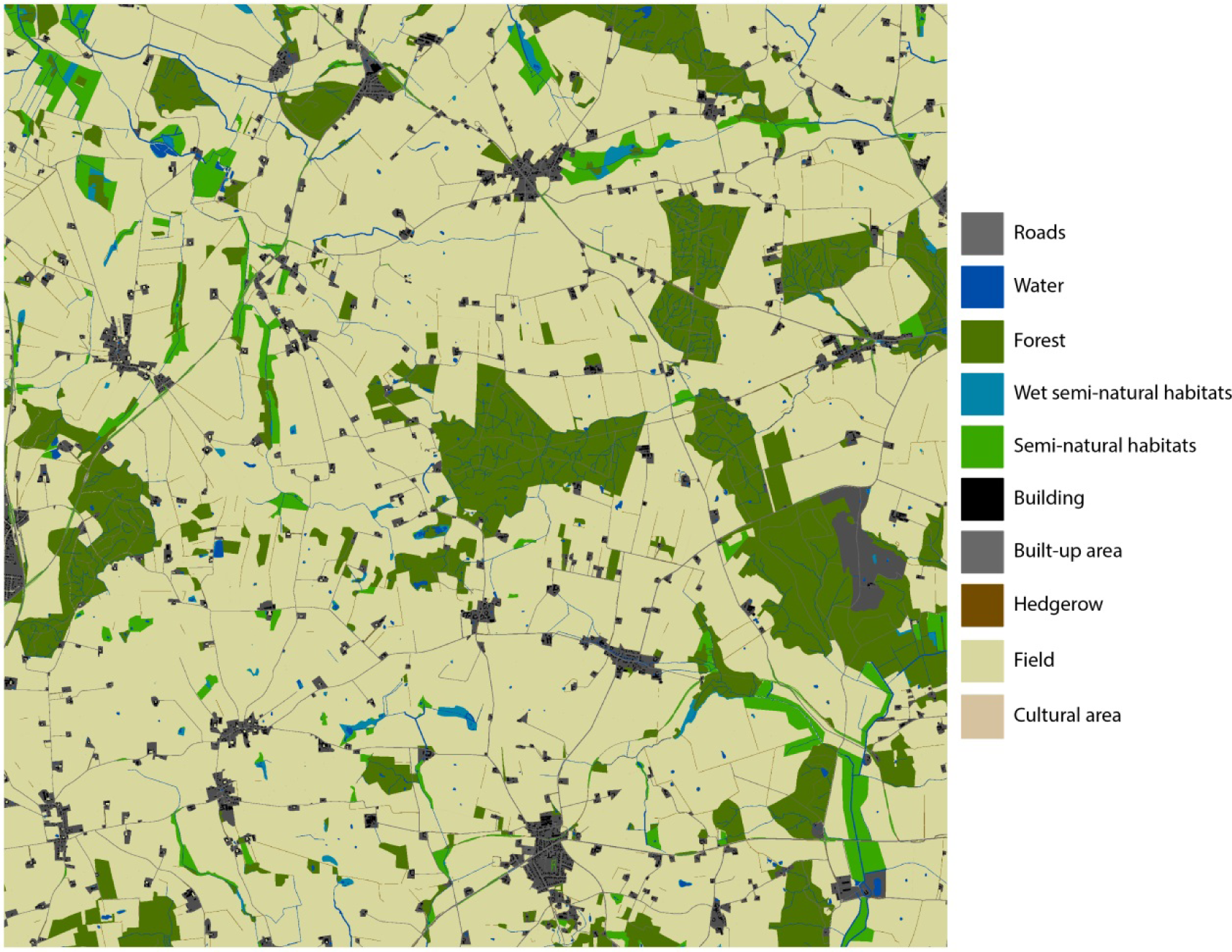
Odder

**Figure 10.**
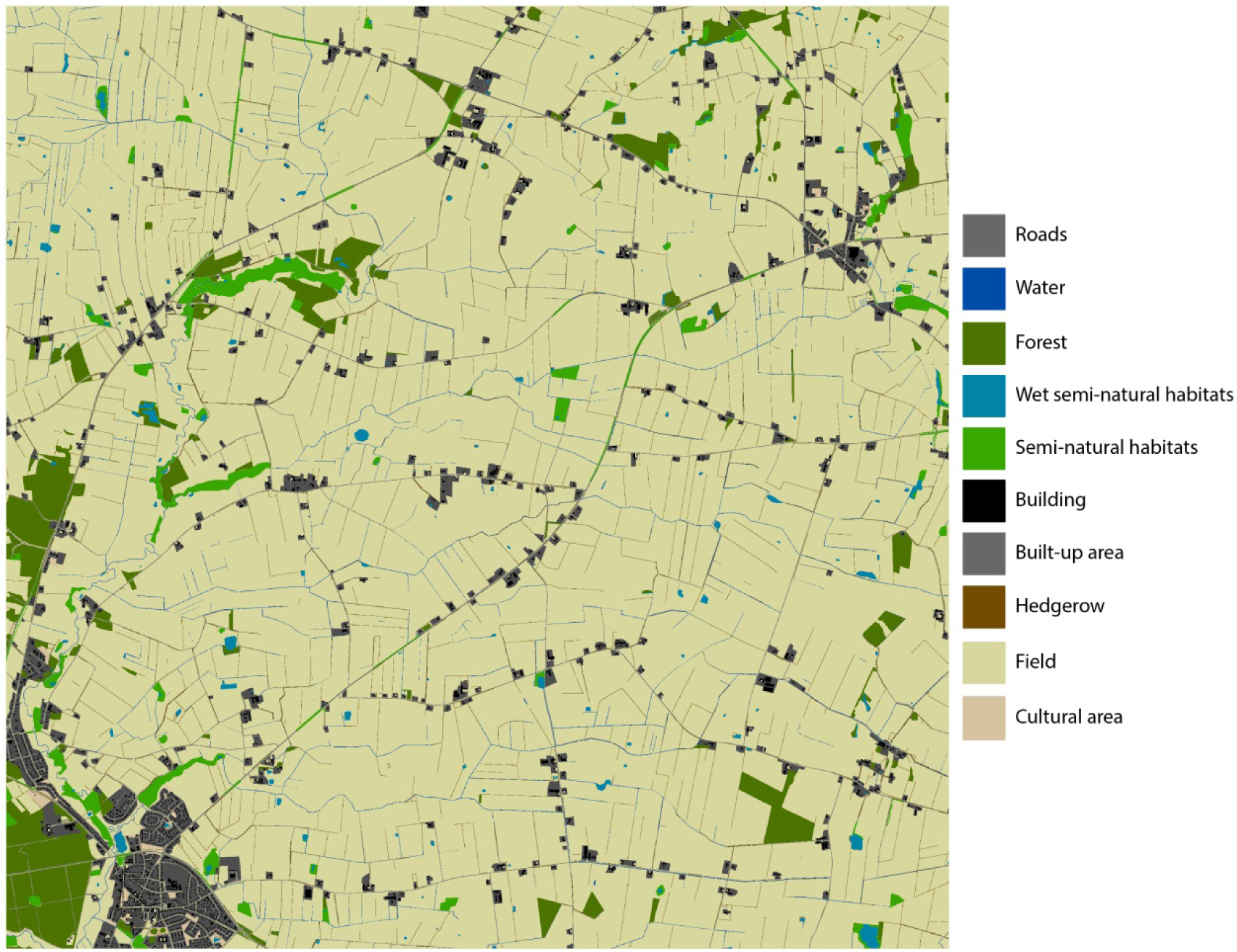
Toftlund

**Figure 11.**
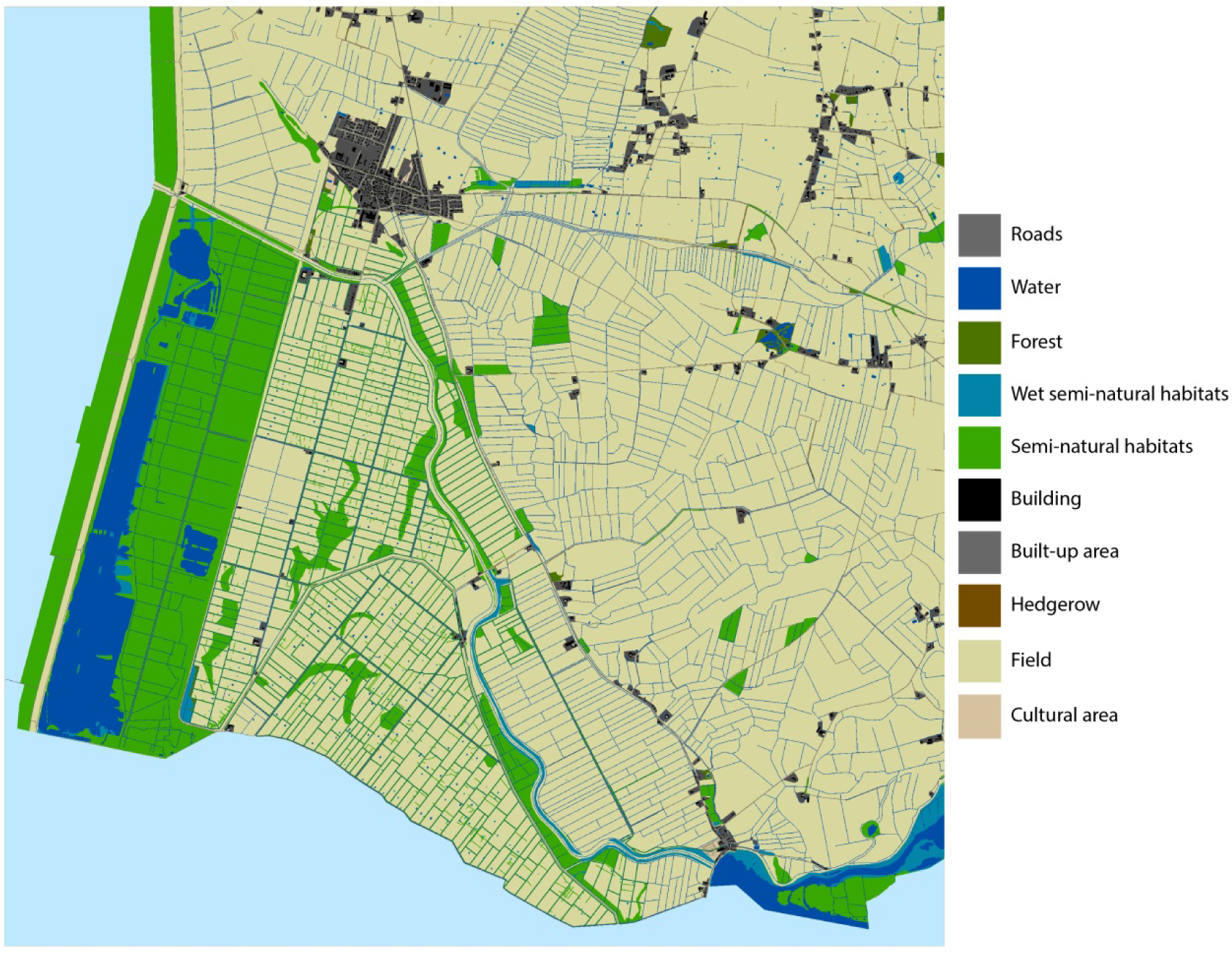
Tønder

## APPENDIX 4: THE PROPORTION OF EACH CROP ASSUMED TO BE GROWN BY EACH FARM TYPE

The proportion of each ALMaSS crop type assumed to be grown by each farm type was based on a national farm classification.

**Table 1:** Conventional farm types and the proportion of each crop or group of crops assumed to be grown.

| Farm Type/Crop | Conv. Pig | Conv. Cattle | Conv. Arable | Conv. Hobby | Conv. Mixed Stock | Conv. Potato | Conv. Beet | Conv. Veg | Other Farm Types | Mean |
| --- | --- | --- | --- | --- | --- | --- | --- | --- | --- | --- |
| Spring Barley | 0.28 | 0.19 | 0.34 | 0.22 | 0.30 | 0.32 | 0.35 | 0.07 | 0.08 | 0.22 |
| Rotational Grass | 0.03 | 0.29 | 0.02 | 0.25 | 0.10 | 0.04 | 0.02 | 0.04 | 0.31 | 0.14 |
| Winter Wheat | 0.32 | 0.07 | 0.32 | 0.09 | 0.19 | 0.07 | 0.28 | 0.02 | 0.02 | 0.11 |
| Permanent Grass | 0.04 | 0.17 | 0.02 | 0.24 | 0.07 | 0.02 | 0.01 | 0.02 | 0.36 | 0.09 |
| Oats | 0.02 | 0.01 | 0.03 | 0.02 | 0.02 | 0.01 | 0.00 | 0.01 | 0.01 | 0.05 |
| Potatoes | 0.00 | 0.00 | 0.00 | 0.00 | 0.00 | 0.18 | 0.00 | 0.03 | 0.02 | 0.05 |
| Winter Rye | 0.04 | 0.01 | 0.04 | 0.02 | 0.04 | 0.03 | 0.00 | 0.00 | 0.00 | 0.04 |
| Sugar Beet | 0.00 | 0.00 | 0.01 | 0.00 | 0.01 | 0.00 | 0.27 | 0.00 | 0.00 | 0.03 |
| Spring Barley Silage | 0.00 | 0.05 | 0.00 | 0.01 | 0.01 | 0.00 | 0.00 | 0.00 | 0.01 | 0.02 |
| Winter Rape | 0.10 | 0.02 | 0.10 | 0.02 | 0.07 | 0.01 | 0.02 | 0.00 | 0.01 | 0.02 |
| Spring Wheat | 0.01 | 0.00 | 0.01 | 0.00 | 0.01 | 0.01 | 0.00 | 0.00 | 0.00 | 0.02 |
| Winter Barley | 0.08 | 0.02 | 0.07 | 0.02 | 0.04 | 0.01 | 0.01 | 0.00 | 0.00 | 0.02 |
| Seed Grass | 0.03 | 0.01 | 0.01 | 0.00 | 0.08 | 0.02 | 0.02 | 0.00 | 0.00 | 0.02 |
| Vegetable crops | 0.00 | 0.00 | 0.00 | 0.00 | 0.00 | 0.00 | 0.00 | 0.22 | 0.00 | 0.02 |
| Maize Silage | 0.01 | 0.13 | 0.00 | 0.01 | 0.02 | 0.03 | 0.01 | 0.00 | 0.01 | 0.02 |
| Others | 0.02 | 0.03 | 0.02 | 0.08 | 0.04 | 0.24 | 0.01 | 0.58 | 0.16 | 0.14 |

**Table 2:**
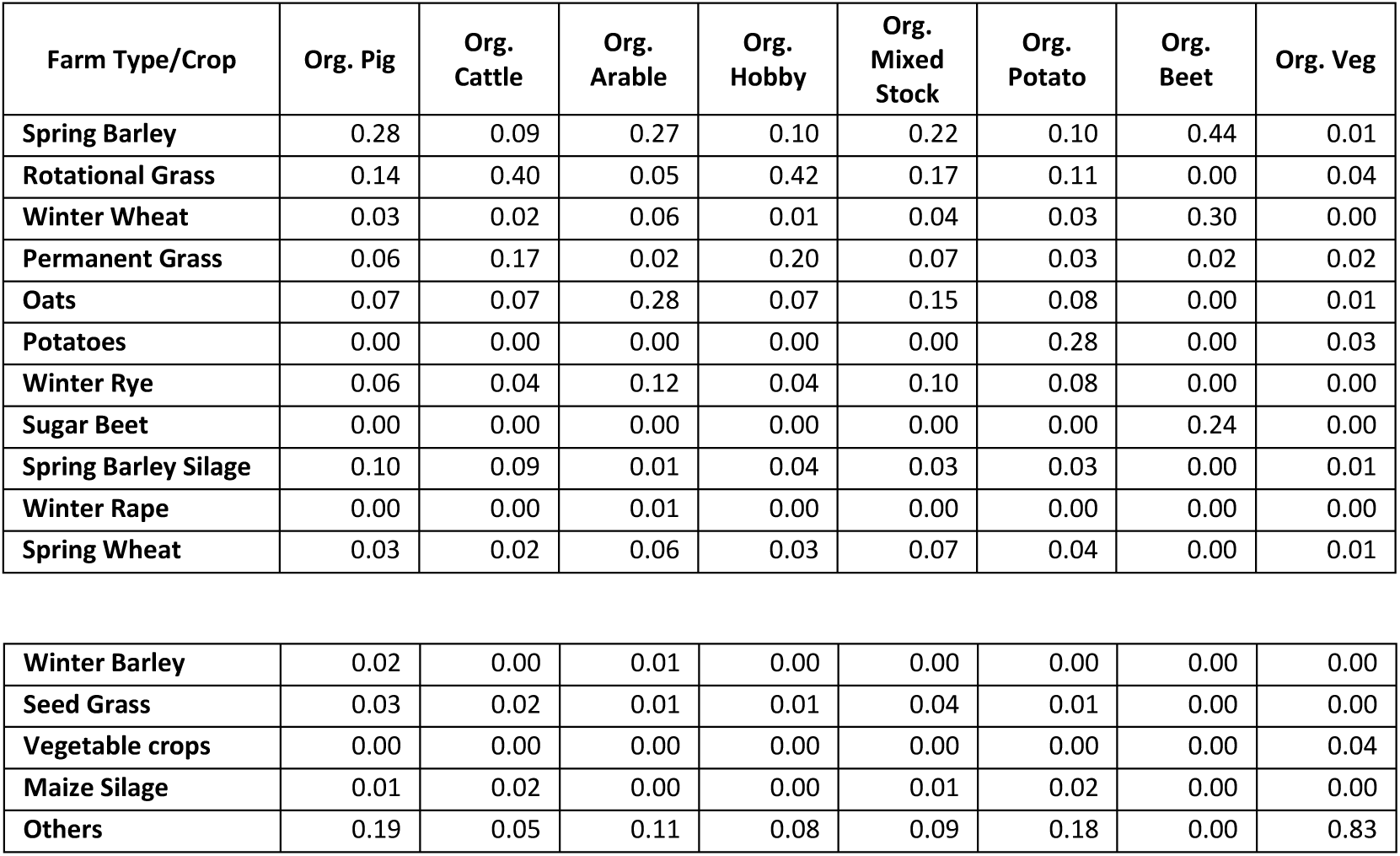
Organic farm types and the proportion of each crop or group of crops assumed to be grown.

